# Discovery of an Achiral Small Molecule TREM2 Agonist with Improved Pharmacokinetic Profile and Validated Target Engagement

**DOI:** 10.1101/2025.06.22.660908

**Authors:** Shaoren Yuan, Farida El Gaamouch, Sungwoo Cho, Katarzyna Kuncewicz, Hossam Nada, Moustafa T. Gabr

## Abstract

The triggering receptor expressed on myeloid cells 2 (TREM2) is a lipid-sensing immunoreceptor on microglia that has emerged as a therapeutic target for Alzheimer’s disease. Here, we report the discovery of **C1**, an achiral structural analog of **VG-3927**—the first small molecule TREM2 agonist to enter clinical development. **C1** was synthesized via a modular and enantioselective-free route using sequential Suzuki couplings, enabling rapid scaffold diversification. Compared to **VG-3927**, the stereochemically simplified derivative (**C1**) exhibits superior microglial phagocytosis and validated target engagement. **C1** induces TREM2 activation in HEK293-hTREM2/DAP12 cells, and its direct binding to TREM2 was confirmed using both microscale thermophoresis (MST) and surface plasmon resonance (SPR). Importantly, **C1** also demonstrates a superior pharmacokinetic profile to **VG-3927**, including enhanced metabolic stability in human and mouse microsomes, favorable PAMPA permeability, and a LogD_7_._4_ compatible with CNS penetration. Docking studies suggested a potential binding mode of **C1** at the extracellular domain of TREM2, revealing key polar and hydrophobic interactions. These findings position **C1** as a synthetically accessible and pharmacokinetically favorable lead for the development of TREM2-targeted therapies

---

Although there is an urgent need to develop effective treatments for Alzheimer’s disease (AD), currently approved drugs do not halt or significantly slow its progression.^1,2^ Characteristic features of AD include the accumulation of amyloid-beta (Aβ) plaques, the formation of neurofibrillary tangles, and the presence of neuroinflammation.^3-5^ The term “neuroinflammation” refers to the inflammatory responses occurring within the central nervous system (CNS).^5^ Microglia, the brain’s resident immune cells, play a central role in this process by producing proinflammatory factors.^6^ Overproduction of these inflammatory molecules contributes to a toxic neural environment that accelerates the progression of AD.^5^ While conventional therapeutic strategies have focused on targets such as cholinergic pathways, β-secretase, Aβ aggregates, and tau pathology, modulating the inflammatory response of microglia has emerged as a promising alternative avenue for AD intervention.^7-9^

Triggering receptor expressed on myeloid cells 2 (TREM2) is a microglia-specific receptor that is predominantly expressed in these CNS immune cells.^10^ TREM2 activates downstream signaling through DAP12—an adaptor protein with an immunoreceptor tyrosine-based activation motif (ITAM) encoded by *TYROBP*—to support key microglial functions, including phagocytosis, cell survival, proliferation, mobility, and lysosomal activity.^10-13^ Since TREM2 mutations were identified as a major genetic risk factor for AD, its role in AD pathogenesis has been extensively studied in preclinical models.^14,15^ In both amyloid mouse models and AD patients carrying the R47H variant, TREM2 deficiency disrupts microglial clustering around Aβ plaques, leading to diffuse plaques and reduced compaction. It also impairs microglial survival during reactive microgliosis, exacerbates Aβ toxicity, and increases neuronal damage.^14,15^ TREM2-deficient microglia fail to adopt a protective transcriptional profile, and while effects on Aβ load vary by model and disease stage, increased neuritic dystrophy is consistently observed.^14,16^ Conversely, overexpressing human TREM2 enhances microglial phagocytosis, boosts related gene expression, and reduces amyloid burden and neuritic damage.^17^ TREM2 agonistic antibodies further support its protective role by promoting plaque compaction, lowering Aβ levels, mitigating neuritic dystrophy, and improving behavioral performance in mouse models.^18,19^

Therapeutic strategies targeting TREM2 have largely focused on antibodies, with limited examples of small molecule approaches reported. Antibody-based therapies for Alzheimer’s disease face several limitations, including poor blood–brain barrier (BBB) penetration, high production costs, potential immunogenicity, and extended half-lives that can lead to prolonged immune-related side effects.^20,21^ In contrast, small molecules offer distinct advantages: they readily cross the BBB, are suitable for oral delivery, carry a lower risk of immunogenicity, and can be more easily optimized for pharmacokinetics (PK), enabling flexible dosing to reduce adverse effects.^22,23^ Additionally, their lower manufacturing costs may improve accessibility.^23^ Incorporating small molecules into AD immunotherapy could significantly advance the field, particularly as combination strategies targeting multiple receptors gain traction. In this context, we recently reported **MG-257** (Fig. 1) as the first small molecule that modulate the interaction between TREM2 and galectin-3.^24^ Remarkably, **VG-3927** (Fig. 1) is the first small molecule TREM2 agonist as a potential therapy for AD.^25^ **VG-3927** is a potent, brain-penetrant TREM2 agonist that induces antiinflammatory microglial activation, reduces AD-related pathology in humanized mice, and shows confirmed CNS activity in primates.^25^

**Fig. 1.**
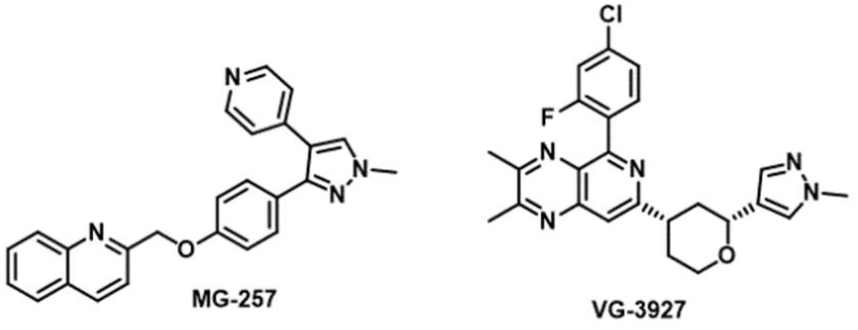
Chemical structures of **MG-257** and **VG-3927**.

As the first small molecule TREM2 agonist to reach clinical development, **VG-3927** establishes a valuable chemical starting point for targeting microglial activation in AD. However, several structural features in **VG-3927** suggest opportunities for medicinal chemistry optimization. The presence of two chiral centers in **VG-3927** adds synthetic complexity and potential enantiomeric liabilities that may complicate scale-up, formulation, and PK profiling. Metabolically labile motifs such as the tetrahydrofuran ring and N-methylated heterocycle may contribute to poor metabolic stability, while its relatively high molecular weight and polar surface area could restrict BBB permeability. Additionally, the halogenated aromatic ring, though potentially beneficial for binding affinity, may carry risks of off-target activity or bioactivation. To address these issues and streamline development, we initiated a design campaign focused on generating achiral derivatives that retain TREM2 agonist activity while improving CNS drug-like properties, metabolic stability, and synthetic accessibility. This approach aims to enable future structure–activity relationship (SAR) development and de-risk the PK and safety profiles of next-generation TREM2 modulators.

The synthetic route shown in Scheme 1 was designed to enable modular access to achiral TREM2 agonist analogs via sequential Suzuki–Miyaura couplings. The synthesis began with the construction of the key heterocyclic scaffold via cross-coupling of an appropriately substituted aryl boronic acid with a 2,3-dichloro-pyrido[3,4-*b*]pyrazine derivative (**A**) under PdCl_2_(dtbpf) catalysis in the presence of cesium carbonate (Cs_2_CO_3_) as base in 1,4-dioxane/water under 60 °C, furnishing the intermediate (**B)**. This step established the central biaryl linkage while retaining a reactive C–Cl handle at the 3-position for subsequent functionalization. Parallel to this, side chain **3** was synthesized via a two-step sequence (Scheme 1) starting from 1-methyl-1*H*-pyrazole-4-carbaldehyde (**1**), followed by the conversion of dihydro-2*H*-pyran-4-yl trifluoromethanesulfonate (**2**) to the corresponding boronic acid pinacol (BPin) ester, enabling its use in the second Suzuki coupling. The final cross-coupling between intermediate **B** and compound **3** was carried out under similar Suzuki conditions, delivering the final set of (**C**) derivatives (Scheme 1). This modular and protecting-group-free route provides efficient access to achiral analogs of the clinical TREM2 agonist, facilitating rapid SAR exploration around both the heterocyclic core and the side chain without the complication of stereocenter control or separation.

**Scheme 1.**
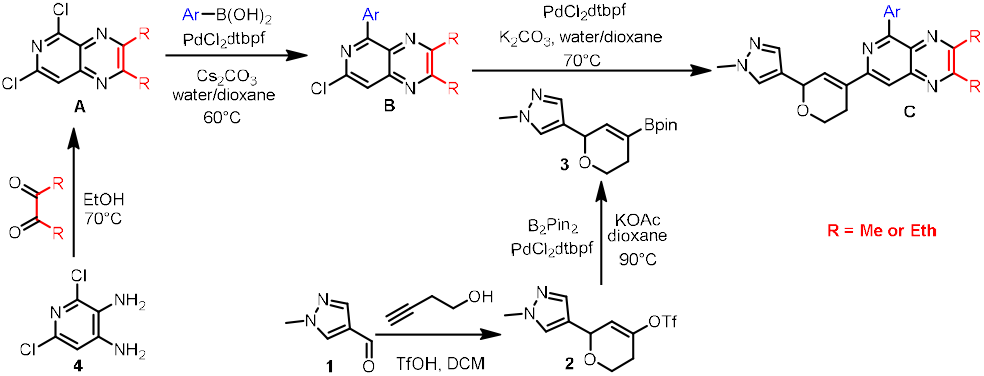
Synthesis of achiral TREM2 agonist analogs (**C**) via sequential Suzuki–Miyaura couplings.

The chemical structures of the **C** derivatives synthesized via the modular route described in Scheme 1 are displayed in Fig. 2. Structural modifications were introduced to probe the contribution of electronic, steric, and solubilizing features to agonist activity, while maintaining the key pharmacophoric elements required for TREM2 engagement. Structural diversification at the C2-position of the pyrido[3,4-b]pyrazine core was achieved by incorporating monosubstituted phenyl rings, as well as extended aromatic systems such as biphenyl and 2-thienyl moieties. Second, the core itself was modified by replacing the methyl substituents with ethyl substituents, introducing a steric and lipophilic shift that may affect receptor binding or metabolic stability. This design enabled exploration of both electronic and spatial parameters within a synthetically accessible, stereochemically simplified scaffold.

**Fig. 2.**
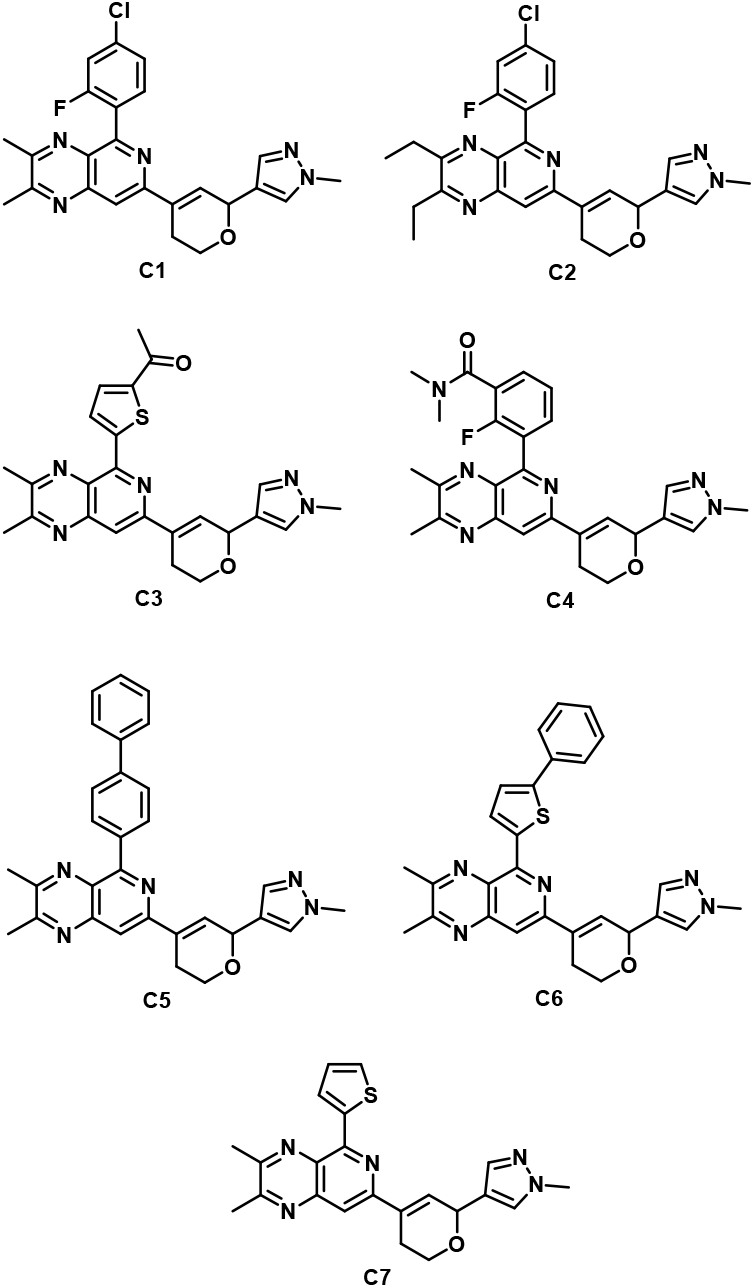
Chemical structures of the synthesized **C1-C7** compounds.

We evaluated **C1**-**C7** compounds for TREM2 pathway activation in HEK293 cells stably expressing human TREM2 and its adaptor protein DAP12 (HEK293-hTREM2/DAP12). Activation of this signaling axis promotes phosphorylation of spleen tyrosine kinase (SYK) at Tyr525/526, a critical event that propagates downstream mechanisms including microglial phagocytosis, survival, and anti-inflammatory responses. Using an AlphaLISA-based assay, SYK phosphorylation was measured following treatment with increasing concentrations of each tested compound, and signal intensity was quantified using a SPARK plate reader (Tecan) under AlphaLISA detection settings. **VG-3927** was used as a control in this study. As shown in Fig. 3A, **VG-3927, C1**, and **C2** induced statistically significant increases in SYK phosphorylation, indicating potent activation of the TREM2–DAP12 signaling cascade. Unlike **VG-3927**, dose-dependent patterns of TREM2 activation were more evident in both **C1** and **C2** compounds. Notably, compound **C5** elicited a slight increase in phospho-SYK levels, however, the magnitude of response was lower than that of **VG-3927**. Other analogs (**C3, C4, C6**, and **C7**) did not produce notable increases in SYK phosphorylation relative to the DMSO control, highlighting the significance of the 2-fluoro-4-chloro phenyl substituent in **C1** and **C2** in driving receptor-mediated signaling. These data establish both **C1** and **C2** as promising candidates for further mechanistic studies of TREM2 agonism.

**Fig. 3.**
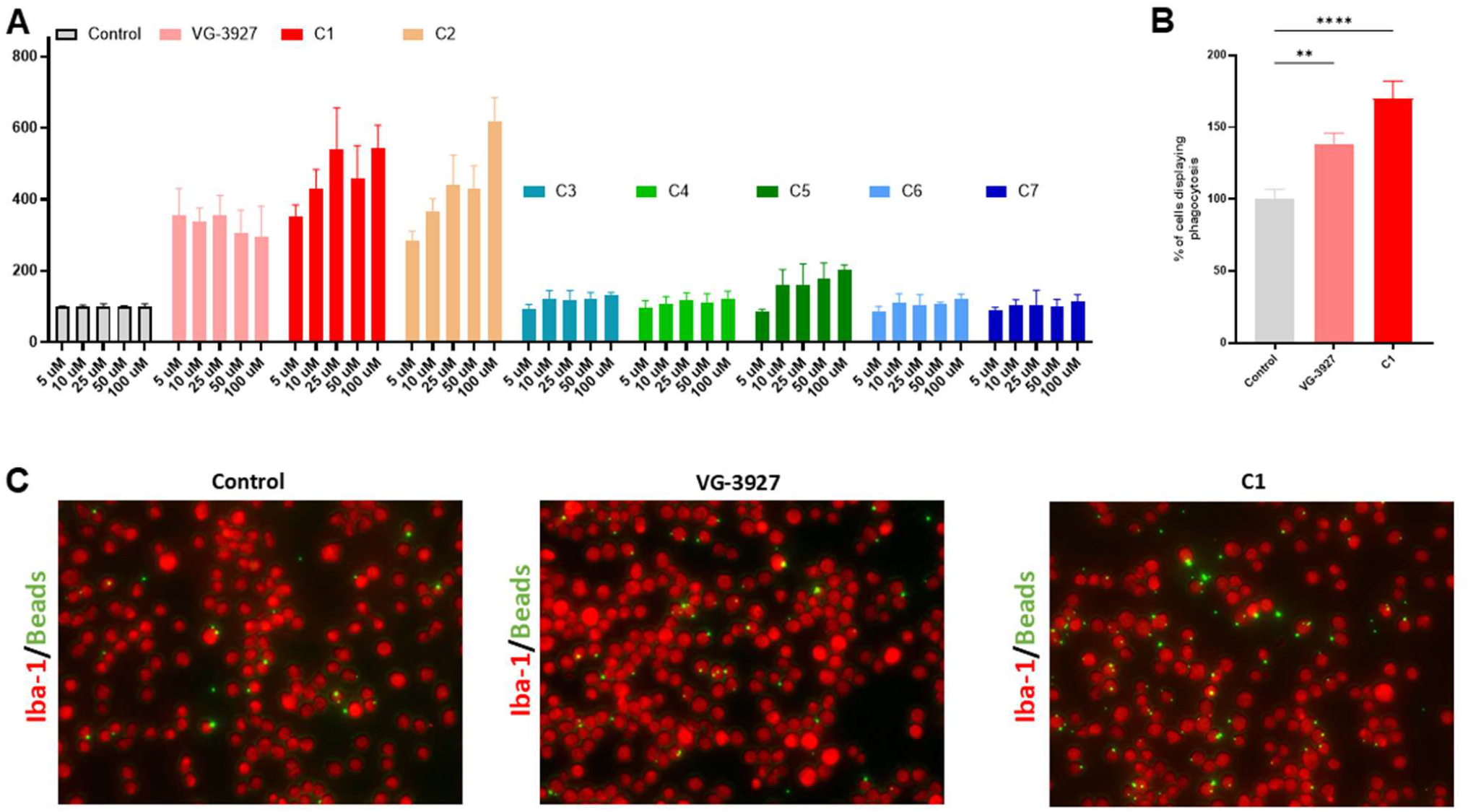
In vitro validation of the TREM2 binding activity. **(A)** Histogram representing the quantification of phospho-Syk levels, measured using AlphaLisa technique, in untreated condition or after treatment with increasing concentrations of **VG-3927** and **C1-C7** compounds. Error bars represent means ± SEM (N=4). (**B)** Histogram representing the quantification of BV2 cells showing at least one fluorescent latex bead in their cell body (percentage of control) after treatment with compound **VG-3927** and **C1** (25 μM-1 h) (n=7-10 pictures/group (N=4)). Statistical analyses were performed using One-Way ANOVA followed by a Kruskal-Wallis multiple comparisons test was done. Error bars represent means ± SEM. \*\**p* < 0.005, \*\*\*\**p* < 0.0001. (**C)** Representative images of BV2 cells immuno-labelled with antibody directed against IBA1 (red) and containing fluorescent beads (green) in their cell body (scale bar=20 μM), in control condition or following treatment with compound **VG-3927** or **C1**.

Glial phagocytosis serves as a vital protective mechanism in various neurodegenerative diseases. As the brain’s innate immune cells, glial cells act as environmental sensors and cleaners, effectively removing cellular debris and damaged components. We explored whether compound **C1** could modulate the phagocytic activity of microglial cells compared to **VG-3927**, potentially influencing neurodegenerative disease progression. To assess phagocytic function, we employed BV2 cells, a murine microglial cell line, and quantified bead uptake using fluorescent latex beads. BV2 cells were serum-deprived prior to treatment with either compound **C1** or **VG-3927**, followed by bead addition. Treatment with compound **C1** enhanced phagocytosis by approximately 1.7 times compared to the control. Similarly, **VG-3927** also enhanced bead uptake, reaching around 1.4 times the control level (Fig. 3B and 3C). Despite both compounds promoting glial phagocytosis, the effect of compound **C1** was significantly greater than that of **VG-3927** (Fig. 3B).

To directly evaluate the molecular interaction between the top-performing compounds **C1** and **C2** and the TREM2 receptor, microscale thermophoresis (MST) was employed as a biophysical technique. Recombinant human TREM2 was fluorescently labeled and incubated with increasing concentrations of the test compounds to monitor concentration-dependent changes in thermophoretic mobility, indicative of direct binding. The results of this MST analysis are summarized in Table 1. Notably, **C1** exhibited a binding profile with a measurable equilibrium dissociation constant (*K*_*D*_) of 71.36 ± 4.06 µM (Fig. 4A), confirming its direct interaction with the TREM2 receptor. In contrast, **C2** and the reference compound **VG-3927** showed minimal or no detectable binding under the same conditions (Table 1). The lack of measurable binding for **VG-3927** is consistent with prior reports suggesting that its agonist activity may arise from TREM2 clustering rather than direct ligand–receptor interaction.^25^ This mechanism could also explain the cellular activity observed for **C2**, which may act through a similar clustering-based pathway. To validate the MST findings and further confirm direct binding of **C1** to TREM2, surface plasmon resonance (SPR) was performed. As shown in Fig. 4B, **C1** displayed dose-dependent binding to immobilized TREM2, as evidenced by the single-cycle kinetic sensorgram. Given the therapeutic advantages of a small molecule that directly engages its molecular target—including improved specificity, predictable SAR, and reduced off-target effects—we prioritized **C1** for further investigation.

**Table 1.** MST analysis of TREM2 binding for **C1, C2**, and **VG-3927**.

| Compound | $K_D$ (μM) |
| --- | --- |
| **C1** | $71.36 \pm 4.06$ |
| **C2** | ND<sup>a</sup> |
| **VG-3927** | ND<sup>a</sup> |

<sup>a</sup>not determined due to negligible impact on $F_{\text{norm}}$ .

**Fig. 4.**
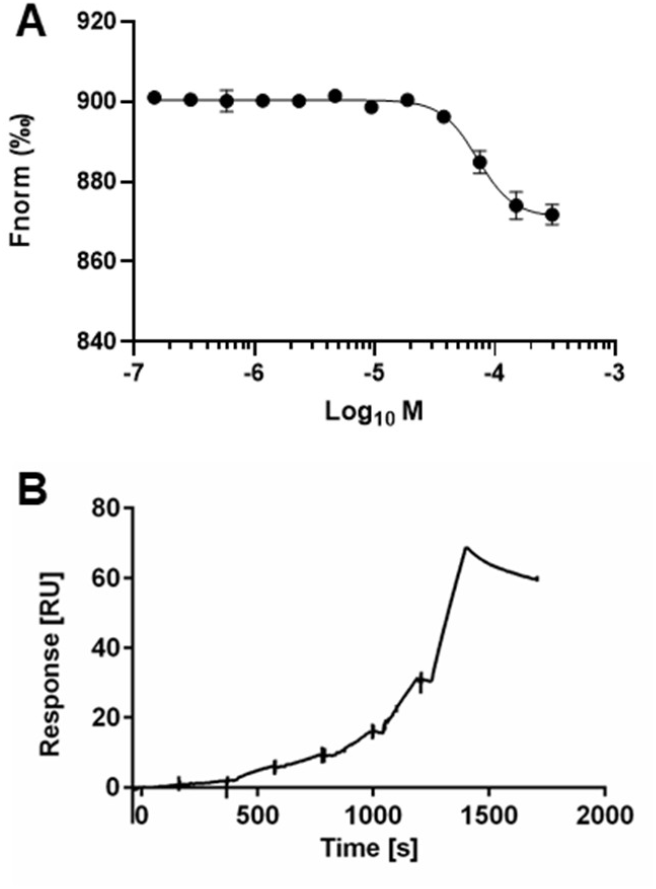
**(A)** Dose-dependent binding experiments for **C1** using MST. Data are presented as Mean ± SEM (n = 3). **(B)** Single-cycle kinetic sensorgram of **C1** interacting with TREM2 using SPR. A two-fold dilution series of **C1**, consisting of seven concentrations ranging between 200 µM and 3.12 µM, were prepared in running buffer (PBS-P supplemented 2% DMSO) and injected over the prepared surface of the SA Sensor Chip.

Given the central role of PK in determining the success of CNS-targeted agents, early ADME profiling is critical for selecting candidates with the best potential for brain exposure, metabolic stability, and safety. The comparative in vitro ADME analysis of **VG-3927** and **C1** (Table 2) provides valuable insights into their drug-like properties and supports the prioritization of **C1** as a more promising therapeutic candidate for AD. While both compounds exhibit favorable solubility and safety profiles, distinct differences in PK parameters highlight the superior characteristics of **C1**. As shown in Table 2, **C1** demonstrates a more favorable balance of lipophilicity and polarity, as reflected by its LogD_7_._4_ value (2.79 versus 2.48 for **VG-3927**) and lower topological polar surface area (TPSA, 68 Å^2^ versus 82 Å^2^). These properties suggest enhanced membrane permeability, a finding supported by the higher Parallel Artificial Membrane Permeability Assay (PAMPA) permeability of **C1** (11.8 × 10?^6^ cm/s) compared to **VG-3927** (8.6 × 10?^6^ cm/s). Such features are particularly desirable in CNS-targeted drug development, where passive diffusion across the blood-brain barrier is often required.

**Table 2.** In vitro PK properties for **VG-3927** and **C1**.

| PK Parameter | VG-3927 | C1 |
| --- | --- | --- |
| $\text{LogD}_{7.4}^a$ | 2.48 | 2.79 |
| TPSA | 82 $\text{\AA}^2$ | 68 $\text{\AA}^2$ |
| **Kinetic Solubility 1%**<br>**DMSO/PBS [μM]<sup>b</sup>** | >100 | >100 |
| **t<sub>1/2</sub> [min] mouse liver micro-**<br>**somes<sup>c</sup> / CL<sub>int</sub><sup>d</sup>** | 31.5 ± 4.2 /<br>42 ± 5.6 | 39.2 ± 5.1 /<br>35 ± 2.3 |
| **t<sub>1/2</sub> [min] human liver mi-**<br>**crosomes<sup>c</sup> / CL<sub>int</sub><sup>d</sup>** | 38.1 ± 3.2 /<br>36.4 ± 2.4 | 48.7 ± 7.1 /<br>27 ± 3.4 |
| **t<sub>1/2</sub> [h] mouse plasma** | 1.5 ± 0.2 | 1.9 ± 0.3 |
| **t<sub>1/2</sub> [h] human plasma** | 1.8 ± 0.4 | 2.4 ± 0.2 |
| **PAMPA permeability**<br>**(cm/s)** | 8.6 × 10<sup>-6</sup> | 11.8 × 10<sup>-6</sup> |
| **PPB human [%]<sup>c</sup>** | 91.4 ± 0.51 | 95.8 ± 0.92 |
| **IC<sub>50</sub> against hERG**<br>**[μM]** | >100 | >100 |

<sup>a</sup>LogD (pH = 7.4) the distribution coefficient at physiological pH 7.4. <sup>b</sup>Kinetic solubility measured in 1% DMSO/PBS (pH 7.4) by UV-vis spectrophotometry (n=3). <sup>c</sup>Values are mean ± SD (n=3). <sup>d</sup>Intrinsic clearance expressed in μL/(min × mg) protein. Values are mean ± SD (n=3).

Metabolic stability, as measured by microsomal half-life and intrinsic clearance, also favors **C1**. In both mouse and human liver microsomes, **C1** exhibited longer half-lives and lower intrinsic clearance rates, indicating reduced susceptibility to metabolic degradation. Specifically, human liver microsomal t_1_/_2_ for **C1** was 48.7 ± 7.1 min versus 38.1 ± 3.2 min for **VG-3927**, with a corresponding lower CL_int_ (27 ± 3.4 versus 36.4 ± 2.4 μL/(min × mg)). These findings suggest that **C1** may have a longer systemic exposure and reduced hepatic clearance in vivo, contributing to improved PK. Plasma stability and protein binding data further underscore favorable ADME profile of **C1** as revealed by exhibiting a longer half-life in both mouse (1.9 ± 0.3 h) and human plasma (2.4 ± 0.2 h), and while it showed slightly higher human plasma protein binding (95.8% vs. 91.4%), this is within an acceptable range and not expected to adversely impact bioavailability or CNS penetration (Table 2). Notably, both compounds showed minimal risk of cardiotoxicity, with hERG IC_50_ values exceeding 100 µM. This, combined with their high solubility (>100 µM) in aqueous buffer, suggests both candidates possess favorable physicochemical properties. However, the superior permeability, metabolic stability, and overall ADME balance observed with **C1** establish it as the more viable candidate.

The structural features of **VG-3927**, specifically the presence of a chiral tetrahydropyran ring, may contribute to its slightly diminished PK characteristics. While conformational restriction and added polarity from such moieties can improve specificity and reduce off-target effects, they may also reduce membrane permeability and introduce unfavorable PK traits, as observed here. In conclusion, the ADME profile of **C1** supports its further advancement as a lead candidate for AD therapy. Its optimal combination of lipophilicity, metabolic stability, and permeability make it more likely to achieve sufficient brain exposure and therapeutic efficacy in vivo.

To facilitate future optimization of **C1** as a TREM2-targeting molecule, we aimed to computationally investigate its binding interactions within the TREM2 binding site. The crystal structure of TREM2 (PDB ID: 5ELI)^26^ was retrieved from the Protein Data Bank. Protein preparation was carried out using Maestro Schrödinger (version 2021.2), and potential small molecule binding pockets were identified using the machine learning-based PrankWeb server.^27,28^ Compound **C1** was docked into the predicted binding site (Fig. 5A) using the XP mode of the Glide module. The docking results revealed that **C1** forms a hydrogen bond between the oxygen atom of its dihydropyran moiety and the Thr82 residue. Additionally, multiple carbon–hydrogen bonds and π–π stacking interactions (Fig. 5B and 5C) were observed between **C1** and surrounding residues, contributing to the stabilization of the binding pose.

**Fig. 5.**
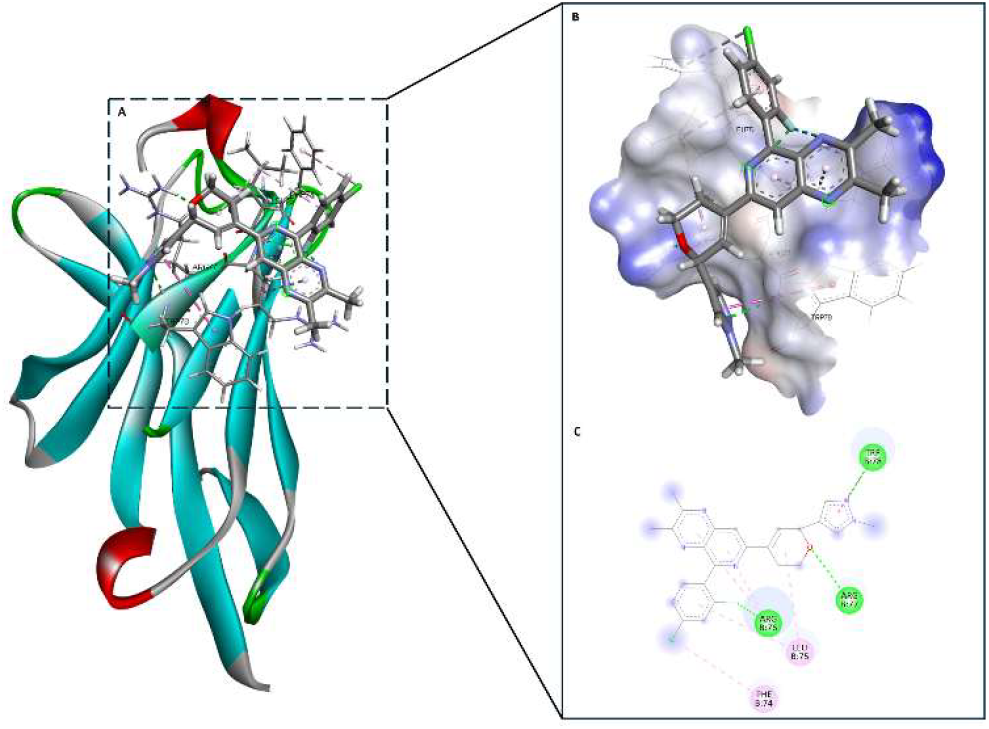
**(A)** Docked complex of **C1** with TREM2. **(B)** 3D model of TREM2 complexed with **C1** based on the crystal structure of TREM2. **(C)** 2D interaction pattern of C1 with the TREM2 predicted active site. Favorable interactions are color-coded as follows: green, hydrogen bonds; π–charge; dark pink, π–π stacking interactions; light green, carbon-hydrogen interactions.

Next, a 100 ns molecular dynamics (MD) simulation was carried out to understand the stability of the predicted **C1**/TREM2 complex. The Root Mean Square Deviation (RMSD) plot of TREM2/**C1** complex (Fig. 6A) shows moderate fluctuations around a mean value close to 2.0 Å which is an indicates that the **C1**/TREM2 complex achieves equilibrium and maintains structural stability throughout the simulation. The RMSD calculations show that after the initial phase of slight deviation, likely corresponding to structural adjustments, the RMSD stabilized which further supports the reliability of the observed dynamics. Subsequently, the stability of the protein/ligand complex was further assessed using DESMOND’s MMGBSA script. The free energy calculations (Fig. 6B) showed that even though the ΔG_bind values fluctuate considerably over time, the ΔG_bind values remained in the negative range which further supports the binding mode predictions. Notably, several deep minima reach below −40 kcal/mol were observed, indicating transient but strong binding conformations. The frequent dips below −20 kcal/mol reflect that the ligand maintains a consistently favorable interaction with TREM2 throughout the simulation. Together, the RMSD and MM/GBSA results suggest that compound **C1** forms a stable and energetically favorable complex with TREM2.

**Fig. 6.**
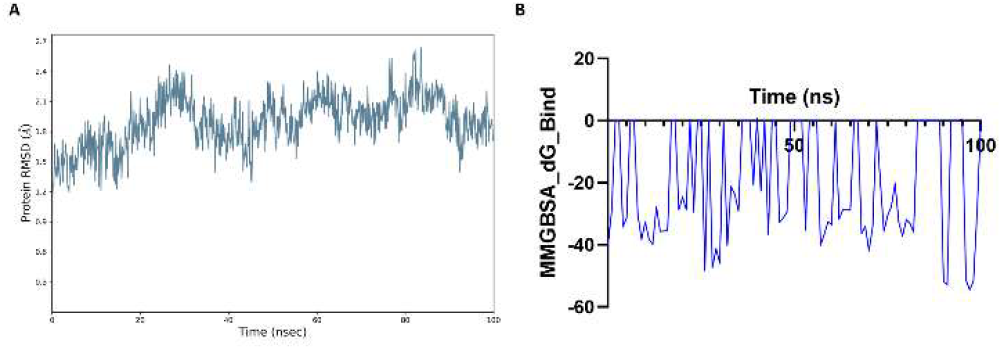
Molecular dynamics analysis of compound C1 bound to TREM2. (A) RMSD of the TREM2/**C1** protein complex over 100 ns, showing system equilibration and structural stability with fluctuations centered around ∼2.0 Å. (B) MM/GBSA-calculated binding free energy (ΔG_bind) of **C1** with TREM2 over 100ns.

In summary, we report the identification of **C1**, an achiral analog of **VG-3927**, with an improved PK profile, retained TREM2 agonist activity, and enhanced microglial phagocytosis. **C1** was synthesized via a modular, enantioselective-free route that enables efficient scaffold diversification. Biophysical assays (MST and SPR) confirmed direct binding of **C1** to TREM2, while computational docking provided structural insights into its binding mode. The compound demonstrates favorable drug-like properties including metabolic stability, PAMPA permeability, and a LogD_7_._4_ supportive of CNS penetration. These results highlight **C1** as a promising lead for further optimization toward TREM2-targeted therapeutics for AD.

## Supporting Information

The Supporting Information is available free of charge on the ACS Publications website.

Experimental procedures, NMR charts, and HRMS data (PDF)

## AUTHOR INFORMATION

### Author Contributions

The manuscript was written through contributions of all authors. All authors have given approval to the final version of the manuscript.

## ABBREVIATIONS

Aβ: amyloid-beta
AD: Alzheimer’s disease
ADME: absorption, distribution, metabolism, excretion
BBB: blood–brain barrier
BPin: boronic acid pinacol ester
CNS: central nervous system
HEK293: human embryonic kidney 293 cells
ITAM: immunoreceptor tyrosine-based activation motif
K_D_: equilibrium dissociation constant
MD: molecular dynamics
MMGBSA: molecular mechanics generalized Born surface area
MST: microscale thermophoresis
PAMPA: Parallel Artificial Membrane Permeability Assay
PK: pharmacokinetics
RMSD: root mean square deviation
SAR: structure–activity relationship
SPR: surface plasmon resonance
TREM2: triggering receptor expressed on myeloid cells 2
TPSA: topological polar surface area
TYROBP: TYRO protein tyrosine kinase binding protein.

## NOTES

The authors declare no competing financial interests.

## ACKNOWLEDGMENTS

This work was supported by the National Institutes on Aging under grant number R01AG083512 (PI: Gabr).

## Insert Table of Contents artwork here

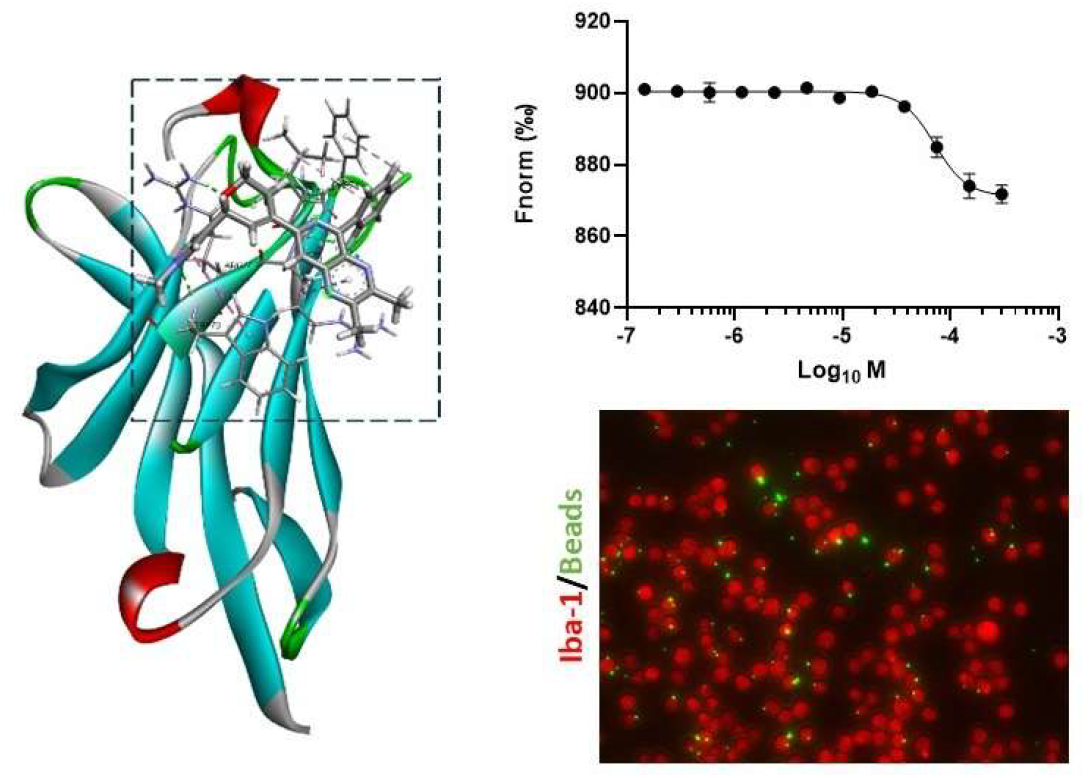

