## Supporting Information for "Discovery of an Achiral Small Molecule TREM2 Agonist with Improved Pharmacokinetic Profile and Validated Target Engagement"

*Electronic Supplementary Information*

**Experimental Procedures**

**1. Chemistry**

All reactions were performed under protection of N_2_ in oven-dried glassware unless water was applied as solvent. To obtain degassed solvent, a continuous flow of N_2_ was bubbled through the solvent for 3 hours, and a N_2_-filled balloon was attached to maintain an inert atmosphere during solvent withdrawal. The degassed solvent can be stored under nitrogen for extended periods without the need for further purging. Chromatographic purification was performed as flash chromatography with Combi-Flash^®^ Rf+ UV-VIS MS COMP using RediSep^®^ Silver normal phase silica gel columns and solvents indicated as eluent with default pressure. Fractions were collected based on UV absorption at 254 nm and/or 280 nm. For compounds lacking UV activity, all eluted fractions were systematically collected and subsequently analyzed by thin-layer chromatography (TLC) using stain visualization. Analytical TLC was performed on Whatman^®^ TLC silica gel UV254 (250 µm) TLC aluminum plates. Visualization was usually accomplished with UV light (254 nm) unless TLC stain was indicated. Proton, carbon and fluorine nuclear magnetic resonance spectra (^1^H NMR, ^13^C NMR and ^19^F NMR) were recorded on a Bruker 500 MHz spectrometer with solvent resonances as the internal standard (^1^H NMR: CDCl_3_ at 7.26 ppm; ^13^C NMR: CDCl_3_ at 77.0 ppm). 1H NMR data are reported as follows: chemical shift (ppm), multiplicity (s = singlet, d = doublet, dd = doublet of doublets, dt = doublet of triplets, ddd = doublet of doublet of doublets, t = triplet, m = multiplet, br = broad), coupling constants (Hz), and integration. The general synthesis of intermediates and final products was adapted from a reported patent.^1^ Mass spectra were obtained through ESI on Micromass Waters LCT Premier XE. The accurate mass analyses run in EI mode were at a mass resolution of 10,000 to 15,000 FWHM and were calibrated using Leucine Enkephalin acetate as an internal standard.

1. **Experimental Procedures**

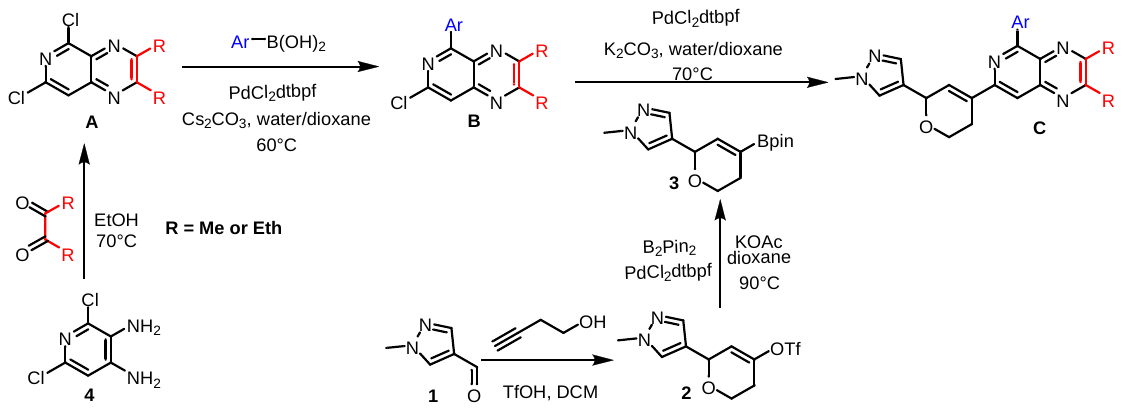

The synthesis of bornic ester side chain intermediate **3:**

Step 1: To a 100 mL dry flask was charged 1-methyl-1*H*-pyrazole-4-carbaldehyde (1.00 g, 9.08 mmol, 1 eq.), which was purged with N_2_. Then 3-butyn-1-ol (955 mg, 1.03 mL, 13.62 mmol, 1.5 eq.) and anhydrous DCM (21 mL) were added. To the vial was added triflic acid (1.64 g, 962 μL, 10.9 mmol, 1.2 eq.) slowly at 0° C. The reaction was warmed to room temperature after 5 min. After 5 h, additional triflic acid (1.64 g, 962 μL, 10.9 mmol, 1.2 eq.) was added. After another 18 h, the mixture was carefully quenched with enough saturated NaHCO_3_ solution and mixture was stirred vigorously until all sticky black tar disappeared. The partitioned mixture was separated, and DCM layer was collected, and the aqueous layer was extracted with DCM (40mL*2). The combined organic layers were washed with brine, dried over Na_2_SO_4_, filtered, and concentrated. The resulting crude material was absorbed onto a plug of silica gel and purified by chromatography (10% −80% EtOAc in hexanes; R_f_ = 0.28 in 70% EtOAc/Hexanes, visualized under anisaldehyde TLC stain) to provide 6-(1-methyl-1*H*-pyrazol-4-yl)-3,6-dihydro-2*H*-pyran-4-yl trifluoromethanesulfonate **2** (1.22 g, 3.91 mmol, 43%) as a light yellow liquid. ^1^H NMR was consistent with the previously reported literature.^1^

Step 2: To a 20 mL scintillation vial was charged 6-(1-methyl-1*H*-pyrazol-4-yl)-3,6-dihydro-2*H*-pyran-4-yltrifluoromethanesulfonate **2** (800 mg, 2.56 mmol, 1 eq.), [1,1′-Bis(di-tert-butylphosphino)ferrocene]dichloropalladium(II) (167 mg, 0.256 mmol, 10 mol%), bis(pinacolato)diboron (976 mg, 3.84 mmol, 1.5 eq.) and potassium acetate (1.01 g, 10.25 mmol, 4 eq.). The flask was purged with N_2_ three times and anhydrous degassed 1,4-dioxane (10.3 mL) was added. The reaction was heated to 90° C for 2 h and the reaction was cooled to room temperature. The reaction mixture was diluted with EtOAc and filtered through a plug of silica gel. The crude material was absorbed onto a plug of silica gel and purified by chromatography (20% −80% EtOAc in hexanes; R_f_ = 0.21 in 70% EtOAc/Hexanes, visualized under anisaldehyde TLC stain) to provide 1-methyl-4-(4-(4,4,5,5-tetramethyl-1,3,2-dioxaborolan-2-yl)-5,6-dihydro-2*H*-pyran-2-yl)-1*H*-pyrazole **3** (498 mg, 1.72 mmol, 67%) as a red oil. ^1^H NMR was consistent with the previously reported literature.^1^

The synthesis of the starting material **A:**

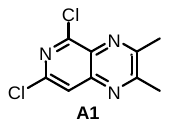
**5,7-dichloro-2,3-dimethylpyrido[3,4-b]pyrazine (A1):** A 100 mL round bottom flask was charged with 3,4-diamino-2,6-dichloropyridine (3 g, 16.9 mmol, 1 eq.) and diacetyl (1.76 mL, 20.2 mmol, 1.2 eq.). EtOH (18 mL) was added to the flask and the mixture was heated to 70 °C. After 5 h, the mixture was filtered through a fritted funnel and the eluent was concentrated to about 9 mL under reduced pressure. Water (9 mL) was added to the solution and the resulting solid was filtered off. The combined solid from both filtrations was washed with hexanes and water twice for each and was allowed to dry on the filter under air to afford 5,7-dichloro-2,3-dimethylpyrido[3,4- b]pyrazine as a light brown solid (3.65g, 16 mmol, 95%). ^1^H NMR was consistent with the previously reported literature.^1^

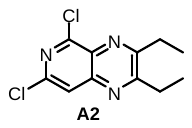
**5,7-dichloro-2,3-diethylpyrido[3,4-b]pyrazine (A2):** A 100 mL round bottom flask was charged with 3,4-diamino-2,6-dichloropyridine (3 g, 16.9 mmol, 1 eq.) and hexane-3,4-dione (2.46 mL, 20.2 mmol, 1.2 eq.). EtOH (15 mL) was added to the flask and the mixture was heated to 70 °C. After 12 h, the solution was concentrated to about 5 mL under reduced pressure, and mixture was filtered through a fritted funnel. Water (5 mL) was added to the filtrate and the resulting solid was filtered off. The combined solid from both filtrations was washed with hexanes, cold methanol and water twice for each and was allowed to dry on the filter under air to afford 5,7-dichloro-2,3-diethylpyrido[3,4- b]pyrazine as a light yellow solid (2.54g, 9.91 mmol, 59%). **^1^H NMR** (500 MHz, CDCl_3_) δ 7.83 (s, 1H), 3.12−3.04 (m, 4H) 1.46 (t, *J* = 7.4 Hz, 3H), 1.42 (t, *J* = 7.4 Hz, 3H). **^13^C NMR** (126 MHz, CDCl_3_) δ 163.90, 160.22, 151.82, 146.30, 145.88, 132.99, 120.83, 28.51, 28.40, 11.44, 11.36. **HRMS(ESI)** m/z: [M-H]^-^ cal. for C11 H10 N3 Cl2, 254.0252; Found, 254.0258.

General method for the synthesis of intermediate **B**: A 20 mL scintillation vial was charged a stir bar, **A** (2 mmol, 1 eq.), corresponding bornic acid ArB(OH)_2_ (2.1 mmol, 1.05 eq.), Cs_2_CO_3_ (1.95 g, 6 mmol, 3 eq.), and [1,1′-Bis(di-tert-butylphosphino)ferrocene] dichloropalladium(II) (65mg, 0.1 mmol, 5 mol%) . A mixture of degassed 1,4-dioxane (5 mL) and water (1.5 mL) was added to the vial. The system was purged with N₂ via three cycles of evacuation and backfilling. The reaction mixture was stirred at 60 °C until the major consumption of starting material (monitored by TLC, anisaldehyde stain was applied when starting materials and products were too close), cooled to room temperature, and partitioned between DCM and water (equal volumes). The organic layer was separated, and the aqueous layer was further extracted with DCM (30 mL). The combined organic layers were washed with brine, dried over anhydrous Na₂SO₄, concentrated under reduced pressure, followed by trituration with cold solution (10 mL, 20% DCM/hexanes). The mixture was vacuum filtered and the solid was washed with same solution (10 mL). Vacuum flow was kept on until the solid was about to dry, then it was washed with cold methanol (2−5 mL) and dried under vacuum to produce intermediate **B**, which can be applied for the next step without further purification.

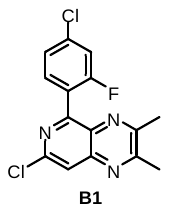
**7-chloro-5-(4-chloro-2-fluorophenyl)-2,3-dimethylpyrido[3,4-b]pyrazine (B1): B1** was obtained (403 mg, 63%) as light brown powder by following the general procedure with **A1** (456 mg) and (4-chloro-2-fluorophenyl)boronic acid (366 mg) under 30 mins reaction time. **B1** was moved forward for the next step without further purification due to only one consolidate spot was observed on TLC (R_f_ = 0.37 with 30% EtOAc/Hexanes).

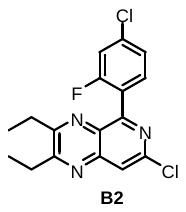
**7-chloro-5-(4-chloro-2-fluorophenyl)-2,3-diethylpyrido[3,4-b]pyrazine (B2):** **B2** was obtained (421 mg, 60%) as light brown powder by following the general procedure with **A2** (512 mg) and (5-acetylthiophen-2-yl)boronic acid (366 mg) under 30 mins reaction time. **B2** was moved forward for the next step without further purification due to only one consolidate spot was observed on TLC (R_f_ = 0.56 with 30% EtOAc/Hexanes).

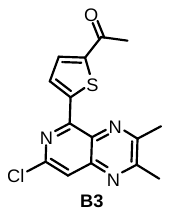
**1-(5-(7-chloro-2,3-dimethylpyrido[3,4-b]pyrazin-5-yl)thiophen-2-yl)ethan-1-one (B3):** **B3** was obtained (251 mg, 40%) as light brown powder by following the general procedure with **A1** (456 mg) and (5-acetylthiophen-2-yl)boronic acid (357 mg) under 12 hours reaction time. **B3** was moved forward for the next step without further purification due to only one consolidate spot was observed on TLC (R_f_ = 0.21 with 30% EtOAc/Hexanes).

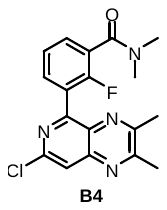
**3-(7-chloro-2,3-dimethylpyrido[3,4-b]pyrazin-5-yl)-2-fluoro-N,N-dimethylbenzamide (B4): B4** was obtained (491 mg, 68%) as light brown powder by following the general procedure with **A1** (456 mg) and (3-(dimethylcarbamoyl)-2-fluorophenyl)boronic acid (443 mg) under 2 hours reaction time. **B4** was moved forward for the next step without further purification due to only one consolidate spot was observed on TLC (R_f_ = 0.22 with 5% MeOH/DCM).

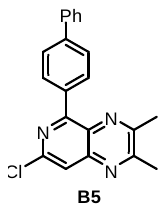
**5-([1,1'-biphenyl]-4-yl)-7-chloro-2,3-dimethylpyrido[3,4-b]pyrazine (B5):** **B5** was obtained (493 mg, 71%) as brown powder by following the general procedure with **A1** (456 mg) and [1,1'-biphenyl]-4-ylboronic acid (416 mg) under 1 hour reaction time. **B5** was moved forward for the next step without further purification due to only one consolidate spot was observed on TLC (R_f_ = 0.32 with 30% EtOAc/Hexanes).

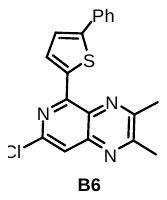
**7-chloro-2,3-dimethyl-5-(5-phenylthiophen-2-yl)pyrido[3,4-b]pyrazine (B6): B6** was obtained (478 mg, 68%) as yellow powder by following the general procedure with **A1** (456 mg) and (5-phenylthiophen-2-yl)boronic acid (428 mg) under 2 hours reaction time. **B6** was moved forward for the next step without further purification due to only one consolidate spot was observed on TLC (R_f_ = 0.32 with 30% EtOAc/Hexanes).

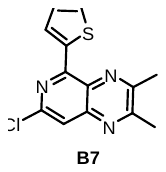
**7-chloro-2,3-dimethyl-5-(thiophen-2-yl)pyrido[3,4-b]pyrazine (B7):** **B7** was obtained (363 mg, 66%) as yellow powder by following the general procedure with **A1** (456 mg) and thiophen-2-ylboronic acid (270 mg) under 3 hours reaction time. **B7** was moved forward for the next step without further purification due to only one consolidate spot was observed on TLC (R_f_ = 0.38 with 30% EtOAc/Hexanes).

General method for the synthesis of final product **C**: A 20 mL scintillation vial was charged a stir bar, **B** (0.5 mmol, 1 eq.), 1-methyl-4-(4-(4,4,5,5-tetramethyl-1,3,2-dioxaborolan-2-yl)-5,6-dihydro-2*H*-pyran-2-yl)-1*H*-pyrazole **3** (131 mg, 0.45 mmol, 0.9 eq.), K_2_CO_3_ (138 mg, 1 mmol, 2 eq.), and [1,1′-Bis(di-tert-butylphosphino)ferrocene] dichloropalladium(II) (49 mg, 75 μmol, 15 mol%). A mixture of degassed 1,4-dioxane (2.25 mL) and water (0.25 mL) was added to the vial. The system was purged with N₂ via three cycles of evacuation and backfilling. The reaction mixture was stirred at 70 °C for 2h, cooled to room temperature, filtered through celite, the pad washed with EtOAc (40 mL) and the filtrate was concentrated under reduced pressure to obtain crude product, which was absorbed onto a plug of silica gel and purified by chromatography. If necessary, trituration with 10 mL solution (5% DCM in hexanes) was applied to the resulting solid to produce the final product **C.**

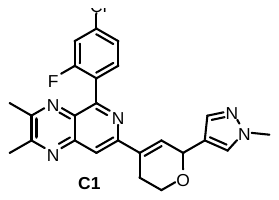
**5-(4-chloro-2-fluorophenyl)-2,3-dimethyl-7-(6-(1-methyl-1*H*-pyrazol-4-yl)-3,6-dihydro-2*H*-pyran-4-yl)pyrido[3,4-b]pyrazine (C1): C1** was obtained (104 mg, 0.231 mmol, 51%) as beige color solid by following general procedure with **3** and **B1** (162 mg), and purification was accomplished by chromatography with 50%−90% EtOAc in hexanes (R_f_ = 0.18 with 100% EtOAc). **^1^H NMR** (500 MHz, CDCl_3_) δ 7.84 (s, 1H), 7.64 (t, *J* = 7.8 Hz, 1H), 7.54 (s, 1H), 7.41 (s, 1H), 7.30 (dd, *J* = 8.3, 2.0 Hz, 1H), 7.24 (dd, *J* = 9.6, 1.9 Hz, 1H), 7.20 – 7.17 (m, 1H), 5.44 (q, *J* = 2.6 Hz, 1H), 4.14 (dt, *J* = 11.3, 4.9 Hz, 1H), 3.96 (td, *J* = 7.1, 3.7 Hz, 1H), 3.89 (s, 3H), 2.84 – 2.78 (m, 1H), 2.76 (s, 3H), 2.75 – 2.70 (m, 1H), 2.68 (s, 3H). **^13^C NMR** (126 MHz, CDCl_3_) δ 160.56 (d, *J* = 253.7 Hz), 158.47, 155.10 (d, *J* = 2.2 Hz), 154.60, 153.05, 144.62, 138.26, 135.79 (d, *J* = 10.3 Hz), 134.52, 133.61, 133.07 (d, *J* = 4.4 Hz), 129.38, 129.15, 125.01, 124.34 (d, *J* = 3.4 Hz), 121.78, 116.42 (d, *J* = 26.0 Hz), 115.22, 68.68, 62.25, 38.93, 25.75, 23.53, 23.38. **^19^F NMR** (471 MHz, CDCl_3_) δ -109.25 (t, *J* = 8.4 Hz). **HRMS (ESI)** m/z: [M+H]^+^ cal. for C24 H22 N5 O F Cl, 450.1497; Found, 450.1516

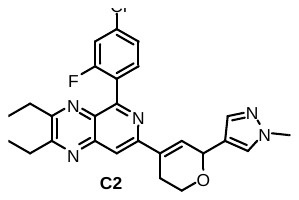

**5-(4-chloro-2-fluorophenyl)-2,3-diethyl-7-(6-(1-methyl-1*H*-pyrazol-4-yl)-3,6-dihydro-2*H*-pyran-4-yl)pyrido[3,4-b]pyrazine (C2): C2** was obtained (103 mg, 0.216 mmol, 48%) as brown color solid by following general procedure with **3** and **B2** (176 mg), and purification was accomplished by chromatography with 30%−80% EtOAc in hexanes (R_f_ = 0.33 with 100% EtOAc). **^1^H NMR** (500 MHz, CDCl_3_) δ 7.88 (s, 1H), 7.70 – 7.65 (m, 1H), 7.54 (s, 1H), 7.41 (s, 1H), 7.29 (dd, *J* = 8.2, 2.1 Hz, 1H), 7.22 (dd, *J* = 9.6, 2.0 Hz, 1H), 7.18 (dd, *J* = 2.7, 1.4 Hz, 1H), 5.44 (q, *J* = 2.7 Hz, 1H), 4.14 (dt, *J* = 11.4, 5.0 Hz, 1H), 3.95 (td, *J* = 7.2, 3.7 Hz, 1H), 3.88 (s, 3H), 3.05 (q, *J* = 7.4 Hz, 2H), 2.99 (q, *J* = 7.3 Hz, 2H), 2.87 – 2.79 (m, 1H), 2.76 – 2.69 (m, 1H), 1.43 (t, *J* = 7.4 Hz, 3H), 1.31 (t, *J* = 7.3 Hz, 3H). **^13^C NMR** (126 MHz, CDCl_3_) δ 162.05, 160.61 (d, *J* = 253.9 Hz), 157.90, 155.15 (d, *J* = 2.3 Hz), 152.94, 144.37, 138.27, 135.59, 134.18, 133.72, 133.09 (d, *J* = 4.4 Hz), 129.10 (d, *J* = 2.4 Hz), 125.16 (d, *J* = 15.4 Hz), 124.17 (d, *J* = 3.4 Hz), 121.79, 116.33, 116.12, 115.46, 68.67, 62.25, 38.89, 28.50, 28.01, 25.79, 11.74, 11.07. **^19^F NMR** (471 MHz, CDCl_3_) δ -108.67 (t, *J* = 8.9 Hz)., **HRMS(ESI)** m/z: [M+H]^+^ cal. for C26 H26 N5 O F Cl, 478.1810; Found, 478.1814.

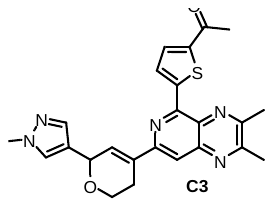
**1-(5-(2,3-dimethyl-7-(6-(1-methyl-1*H*-pyrazol-4-yl)-3,6-dihydro-2*H*-pyran-4-yl)pyrido[3,4-b]pyrazin-5-yl)thiophen-2-yl)ethan-1-one (C3): C3** was obtained (115 mg, 0.258 mmol, 57%) as orange-red color solid by following general procedure with **3** and **B3** (160 mg), and purification was accomplished by chromatography with 60%−100% EtOAc in hexanes (R_f_ = 0.12 with 100% EtOAc). **^1^H NMR** (500 MHz, CDCl3) δ 8.63 (d, J = 4.1 Hz, 1H), 7.73 (d, J = 4.5 Hz, 2H), 7.56 (s, 1H), 7.43 (s, 1H), 7.24 – 7.19 (m, 1H), 5.47 (q, J = 2.8 Hz, 1H), 4.14 (dt, J = 10.6, 5.0 Hz, 1H), 3.96 (ddd, J = 11.6, 7.1, 4.6 Hz, 1H), 3.90 (s, 3H), 2.83 – 2.72 (m, 2H), 2.81 (s, 3H), 2.75 (s, 3H), 2.62 (s, 3H). **^13^C NMR** (126 MHz, CDCl_3_) δ 191.49, 158.51, 154.30, 152.70, 150.21, 149.01, 146.54, 145.14, 138.52, 133.53, 133.12, 132.51, 131.39, 129.49, 129.13, 121.60, 114.92, 68.62, 62.10, 38.95, 26.96, 25.67, 23.45, 23.27. **HRMS(ESI)** m/z: [M+Na]^+^ cal. for C24 H23 N5 O2 S Na, 468.1470; Found, 468.1466.

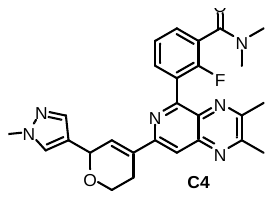
**3-(2,3-dimethyl-7-(6-(1-methyl-1*H*-pyrazol-4-yl)-3,6-dihydro-2*H*-pyran-4-yl)pyrido[3,4-b]pyrazin-5-yl)-2-fluoro-N,N-dimethylbenzamide (C4): C4** was obtained (105 mg, 0.216 mmol, 48%) as orange-yellow color solid by following general procedure except using PdCl_2_dppf•DCM (61 mg) as catalyst with **3** and **B4** (180 mg), and purification was accomplished by chromatography with 2%−5% MeOH in DCM (R_f_ = 0.11 with 5% MeOH/DCM). **^1^H NMR** (500 MHz, CDCl_3_) δ 7.83 (d, *J* = 1.4 Hz, 1H), 7.72 (t, *J* = 7.1 Hz, 1H), 7.56 – 7.51 (m, 2H), 7.40 (s, 1H), 7.35 (td, *J* = 7.6, 1.6 Hz, 1H), 7.20 (s, 1H), 5.43 (s, 1H), 4.13 (dt, *J* = 10.2, 5.1 Hz, 1H), 3.97 – 3.91 (m, 1H), 3.86 (s, 3H), 3.13 (s, 3H), 3.08 (s, 3H), 2.83 – 2.66 (m, 5H), 2.62 (s, 3H). **^13^C NMR** (126 MHz, CDCl_3_) δ 166.75, 158.36, 156.11 (d, *J* = 251.1 Hz), 155.43, 154.36, 153.09, 144.53, 138.39, 134.50, 133.55, 133.29 (d, *J* = 3.6 Hz), 129.92 (d, *J* = 4.1 Hz), 129.40, 129.04, 126.81 (d, *J* = 15.9 Hz), 124.58 (d, *J* = 19.1 Hz), 124.37 (d, *J* = 3.6 Hz), 121.66, 115.20, 68.67, 62.19, 38.88, 38.39 (d, *J* = 3.2 Hz), 35.01, 25.71, 23.49, 23.24.**^19^F NMR** (471 MHz, CDCl_3_) δ -113.64. **HRMS(ESI)** m/z: [M+H]^+^ cal. for C27 H28 N6 O2 F, 487.2258; Found, 487.2239

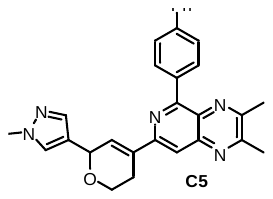
**5-([1,1'-biphenyl]-4-yl)-2,3-dimethyl-7-(6-(1-methyl-1*H*-pyrazol-4-yl)-3,6-dihydro-2*H*-pyran-4-yl)pyrido[3,4-b]pyrazine (C5):** **C5** was obtained (101 mg, 0.213 mmol, 47%) as orange-yellow color solid by following general procedure with **3** and **B5** (174 mg), and purification was accomplished by chromatography with 50%−90% EtOAc in hexanes (R_f_ = 0.14 with 100% EtOAc). **^1^H NMR** (500 MHz, CDCl_3_) δ 8.40 – 8.36 (m, 2H), 7.79 (s, 1H), 7.75 (dd, *J* = 8.4, 1.7 Hz, 2H), 7.72 – 7.67 (m, 2H), 7.57 (s, 1H), 7.48 (dd, *J* = 8.5, 7.0 Hz, 2H), 7.43 (s, 1H), 7.38 (t, *J* = 7.4 Hz, 1H), 7.28 (q, *J* = 1.9 Hz, 1H), 5.47 (q, *J* = 2.7 Hz, 1H), 4.16 (dt, *J* = 10.4, 4.9 Hz, 1H), 3.98 (ddd, *J* = 11.7, 7.4, 4.6 Hz, 1H), 3.89 (s, 3H), 2.86 (dtt, *J* = 14.6, 4.9, 2.3 Hz, 1H), 2.80 – 2.73 (m, 1H), 2.78 (s, 3H), 2.77 (s, 3H). **^13^C NMR** (126 MHz, CDCl_3_) δ 157.70, 157.45, 153.93, 152.67, 145.41, 141.98, 140.87, 138.53, 136.89, 134.28, 133.92, 131.81, 129.12, 129.09, 128.79, 127.47, 127.21, 126.66, 121.80, 114.33, 68.71, 60.35, 38.93, 25.78, 23.45. **HRMS(ESI)** m/z: [M+H]^+^ cal. for C30 H28 N5 O, 477.2294; Found, 474.2309.

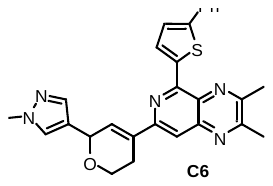
**2,3-dimethyl-7-(6-(1-methyl-1*H*-pyrazol-4-yl)-3,6-dihydro-2*H*-pyran-4-yl)-5-(5-phenylthiophen-2-yl)pyrido[3,4-b]pyrazine (C6):** **C6** was obtained (106 mg, 0.221 mmol, 49%) as orange-red color solid by following general procedure with **3** and **B6** (177 mg), and purification was accomplished by chromatography with 50%−90% EtOAc in hexanes (R_f_ = 0.11 with 100% EtOAc). **^1^H NMR** (500 MHz, CDCl_3_) δ 8.68 (d, *J* = 4.0 Hz, 1H), 7.74 (d, *J* = 7.7 Hz, 2H), 7.64 (s, 1H), 7.59 (s, 1H), 7.45 (s, 1H), 7.44 – 7.38 (m, 3H), 7.32 (t, *J* = 7.4 Hz, 1H), 7.25 (d, *J* = 2.4 Hz, 1H), 5.49 (q, *J* = 2.7 Hz, 1H), 4.15 (dt, *J* = 11.5, 5.1 Hz, 1H), 3.97 (ddd, *J* = 11.6, 7.2, 4.6 Hz, 1H), 3.90 (s, 3H), 2.85 – 2.78 (m, 4H), 2.76 – 2.71 (m, 4H). **^13^C NMR** (126 MHz, CDCl_3_) δ 157.96, 153.62, 152.53, 151.22, 148.53, 145.21, 141.52, 138.60, 134.43, 133.81, 133.02, 132.60, 129.21, 129.00, 128.90, 127.90, 125.98, 124.02, 121.81, 113.45, 68.67, 62.19, 38.96, 25.70, 23.40, 23.26. **HRMS(ESI)** m/z: [M+H]^+^ cal. for C28 H26 N5 O S, 480.1858; Found, 480.1882.

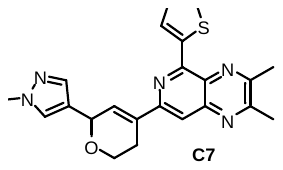
**2,3-dimethyl-7-(6-(1-methyl-1*H*-pyrazol-4-yl)-3,6-dihydro-2*H*-pyran-4-yl)-5-(thiophen-2-yl)pyrido[3,4-b]pyrazine (C7): C7** was obtained (102 mg, 0.251 mmol, 56%) as brown-yellow color solid by following general procedure with **3** and **B7** (138 mg), and purification was accomplished by chromatography with 50%−90% EtOAc in hexanes (R_f_ = 0.15 with 100% EtOAc). **^1^H NMR** (500 MHz, CDCl_3_) δ 8.70 (d, *J* = 3.9 Hz, 1H), 7.65 (s, 1H), 7.58 (s, 1H), 7.52 (d, *J* = 5.1 Hz, 1H), 7.44 (s, 1H), 7.24 (dt, *J* = 3.0, 1.5 Hz, 1H), 7.19 (dd, *J* = 5.1, 3.8 Hz, 1H), 5.47 (q, *J* = 2.7 Hz, 1H), 4.14 (dt, *J* = 10.3, 4.9 Hz, 1H), 3.96 (ddd, *J* = 7.9, 5.4, 2.8 Hz, 1H), 3.89 (s, 3H) 2.84 – 2.77 (m, 4H), 2.76 – 2.72 (m, 4H). **^13^C NMR** (126 MHz, CDCl_3_) δ 157.97, 153.68, 152.50, 151.42, 145.22, 142.12, 138.54, 133.77, 132.84, 131.36, 130.22, 129.12, 129.06, 127.78, 121.81, 113.54, 68.69, 62.24, 38.95, 25.67, 23.39, 23.22. **HRMS(ESI)** m/z: [M+H]^+^ cal. for C22 H22 N5 O S, 404.1545; Found, 404.1551.

**2. MST study for TREM2 binding**

Binding affinity measurements were performed using MST on a Monolith NT.115 system (NanoTemper Technologies, Munich, Germany). TREM2 (BioTechne, MN, USA) was labeled with RED-tris-NTA His-tag labeling kit (NanoTemper Technologies, Munich, Germany) according to manufacturer's instructions. TREM2 assays used the same buffer with 0.005% Tween-20. Labeled protein (40 nM) was incubated with compound at indicated concentrations for 10 minutes at room temperature. Measurements were performed using red filter, 100% LED power, and medium MST power. Data were analyzed with MO. Affinity Analysis software and GraphPad Prism (GraphPad Software, CA, USA) to determine KD values.

**3. SPR for TREM2 binding assessment**

The interaction between the compound and biotinylated human TREM2 (Cat.: 11084-H49H-B, Sino Biological, Beijing, China) was determined by SPR (Biacore 8K, Cytiva). The biotinylated TREM2 protein was immobilized on SA Sensor Chip (Cytiva) in a PBS-P (0.2 M phosphate buffer with 27 mM KCl, 1.37 M NaCl and 0.5% Surfactant P20, pH 7.4, Cytiva) to a response level of 4,343 RU ± 399 RU. Kinetic measurements were run using a single cycle kinetic approach. A two-fold dilution series of the compound, consisting of seven concentrations ranging between 200 µM and 3.12 µM, were prepared in running buffer (PBS-P supplemented 2% DMSO) and injected over the prepared surface of the SA Sensor Chip. Experiments were conducted at 25°C employing the following parameters: flow rate of 30 µl/min, contact time of 200 s, and dissociation time of 900 s. After each analysis an additional wash with 50% DMSO solution was performed. Each interaction was investigated at least in triplicate. The results are presented as sensorgrams obtained after subtraction of the background response signal from a reference flow cell and from a control experiment with buffer injection. The obtained data were analyzed using BiacoreTM Insight Evaluation Software (Cytiva).

**4.** **AlphaLISA pSyk assay**

Phospho-AlphaLISA assay measures a protein target when phosphorylated at a specific residue. The assay uses two antibodies which recognize the phospho epitope and a distal epitope on the targeted protein. In the presence of phosphorylated protein, luminescent Alpha signal is generated. The amount of light emission is directly proportional to the quantity of phosphoprotein present in the sample (from Revvity website). Briefly, HEK –hTREM2/DAP12 cells were seeded at 50,000 cells per well in 96-well plate, in a final volume of 100 μl of growth media composed of DMEM (Gibco) supplemented with 10% FBS (Gibco) and incubated for 24 h at 37 °C, 5% CO_2_. The tested compounds were diluted in media at 5, 10, 25, 50, and 100 μM, growth media was manually removed from the wells and subsequently replaced with the test compounds containing media. Control wells were treated with the vehicle buffer used to solubilize the several compounds. After 1h, the media was gently removed, lysis buffer was added and after complete lysis, alphalisa assay was performed according to manufacturer instructions (Revvity, USA).

**5. Phagocytosis assay**

Phagocytosis assay was performed in BV2 cells using green fluorescent latex beads (Sigma #L1030). Prior use, beads were opsonized in FBS (1 h at 37 °C), the final concentrations for beads and FBS in DMEM were 0.01% (v/v) and 0.05% (v/v) respectively. Briefly, BV2 cells were submitted to serum starvation prior treatment with the tested compound (25 μM for 30 min) or solubilization buffer (control condition) and beads were added to the media for an additional 30 min. Cultures were then washed 3 times with ice-cold PBS and fixed in 4% paraformaldehyde (PFA) (Thermofisher, MA, USA) before undergoing immunocytochemistry. Following Immunostaining, pictures were taken for each condition under 20X objective, total number of cells was counted manually using DAPI staining and each IBA1 immunolabelled-cell containing at least one bead was counted as positive for phagocytosis.

**6.** **Immunocytochemistry**

Fixed cells were permeabilized with 0.25% Triton X-100-containing PBS and subsequently blocked with 1% BSA. Immunocytochemistry was then performed using primary anti IBA1 (Wako, Richmond, VA) antibody followed by secondary 594 Alexa fluorescent antibody (Thermofisher, MA, USA).

**7.** **In vitro PK profiling**

These studies were performed as we previously reported.^2^ The procedures involve the determination of LogD7.4 values, microsomal stability, kinetic solubility, and cytotoxicity against a panel of cell lines. Solubility studies were performed using UV−vis spectrophotometry. PrestoBlue cell viability assay was used to assess the cell viability.

**8. In silico methodology**

The crystal structure of TREM2 (PDB ID: 5ELI) was retrieved from the Protein Data Bank and prepared using Maestro Schrödinger (version 2021.2). Potential small-molecule binding pockets were identified using the machine learning-based PrankWeb server (accessed 05-02-2025). Molecular docking of compound C1 into the predicted binding site was performed using the XP mode of the Glide module to determine the binding pose and interaction patterns. The stability of the predicted C1/TREM2 complex was further assessed through a 100 ns molecular dynamics (MD) simulation. Root Mean Square Deviation (RMSD) values were calculated to evaluate structural stability, and binding free energies were estimated using DESMOND's MMGBSA script to quantify the energetic favorability of the protein-ligand interactions throughout the simulation period. Discovery studio visualizer program was used to display the 3D and 2D complexes while GraphPad software was used to plot the MMGBSA plot.

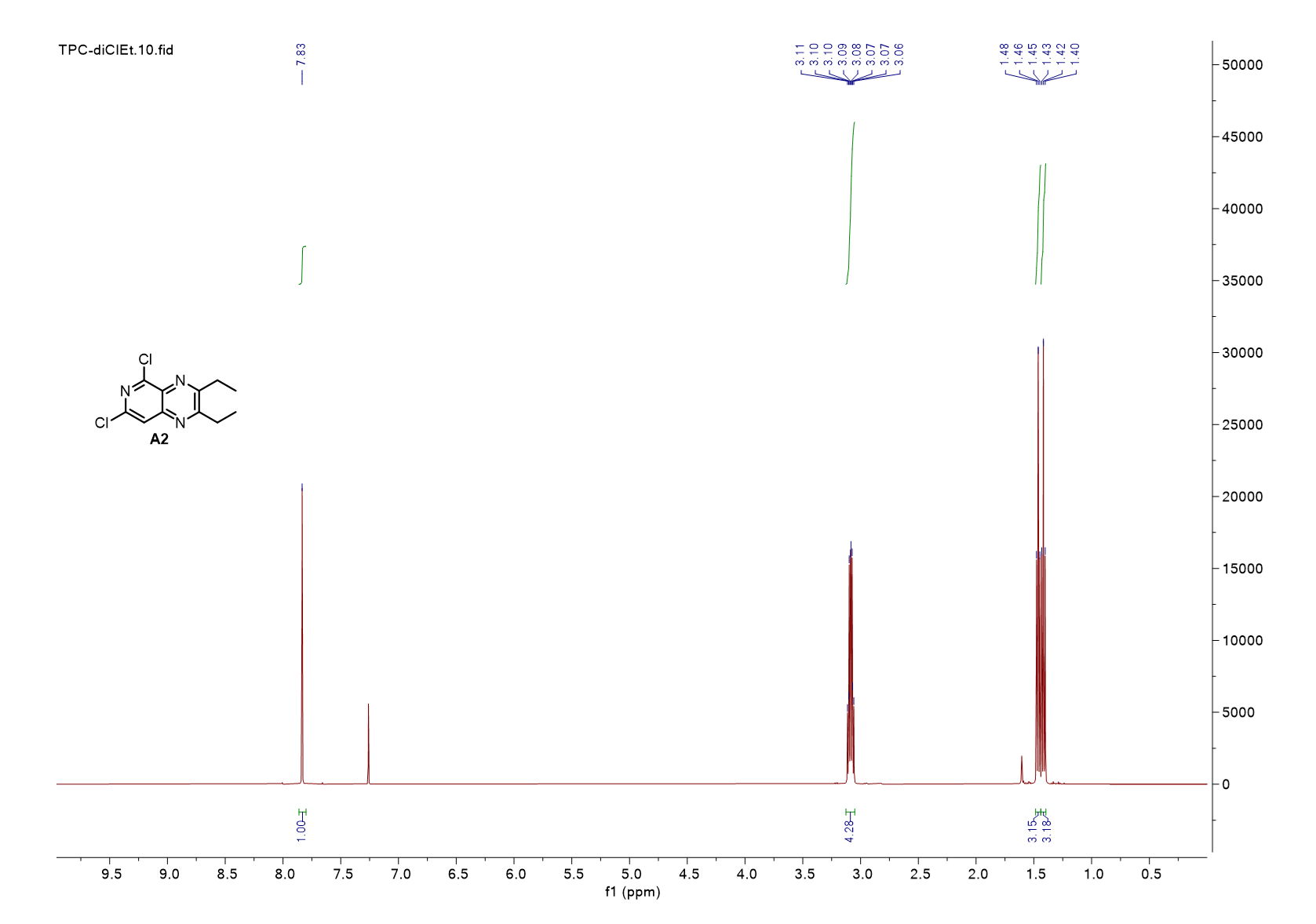

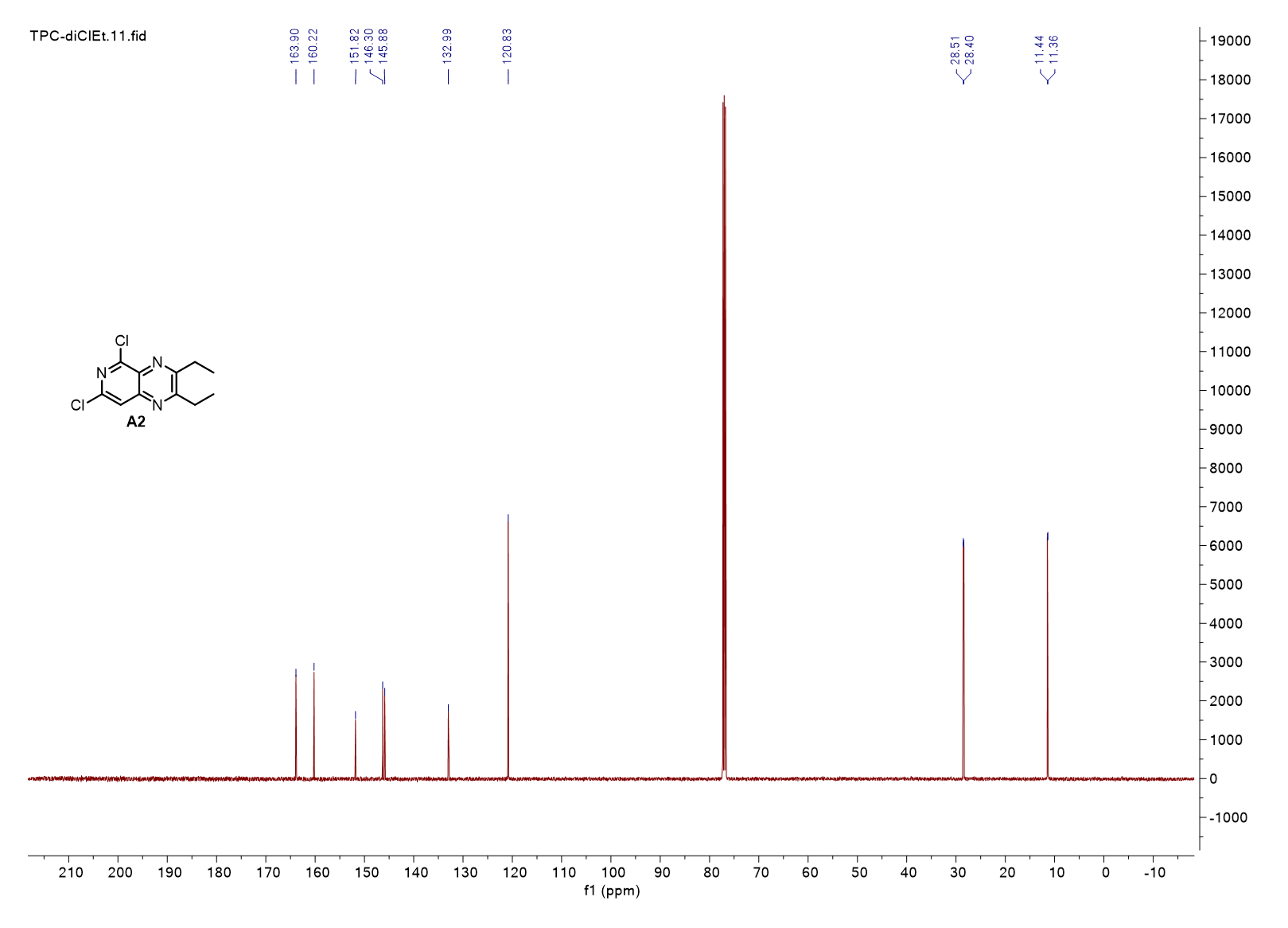

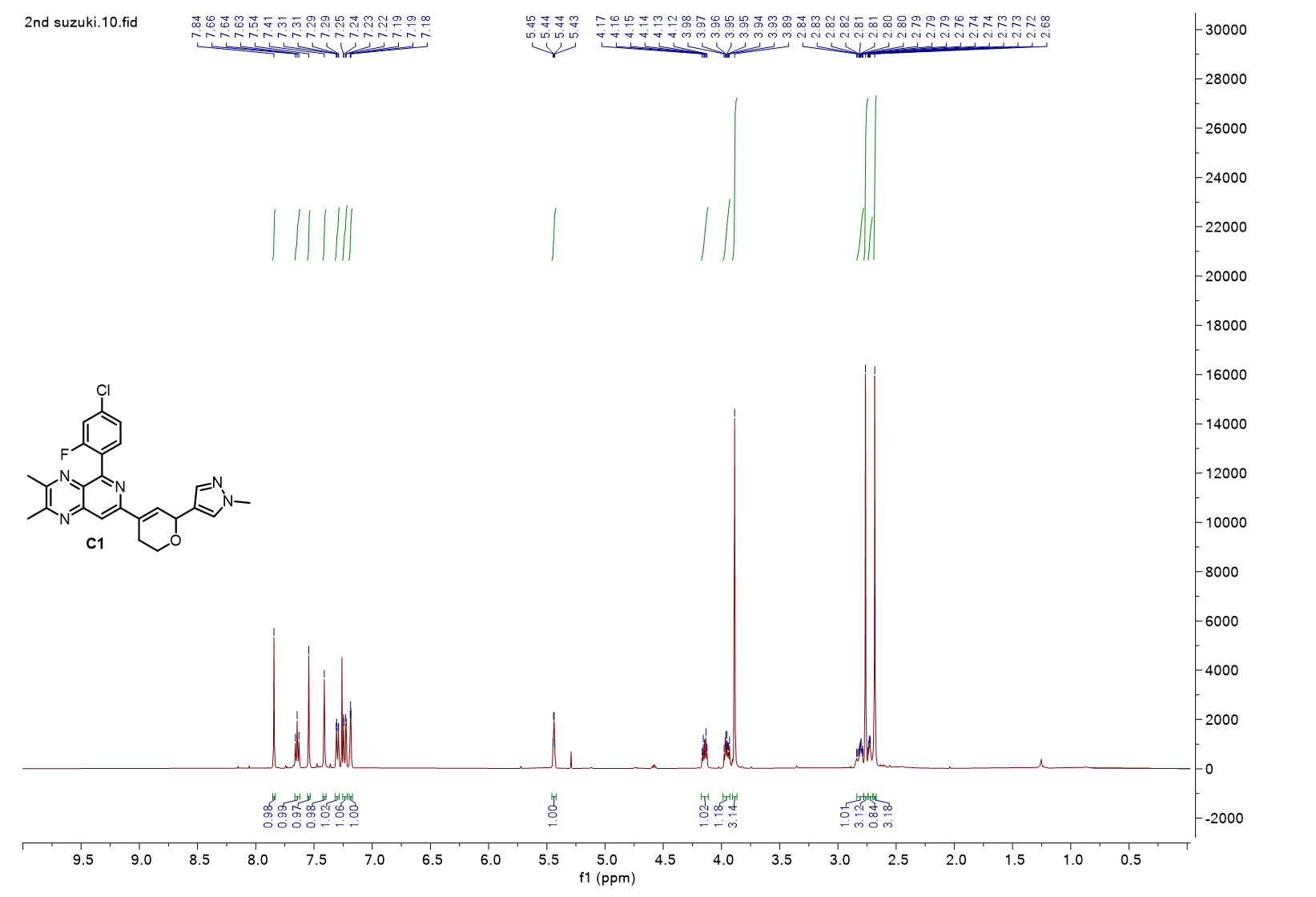

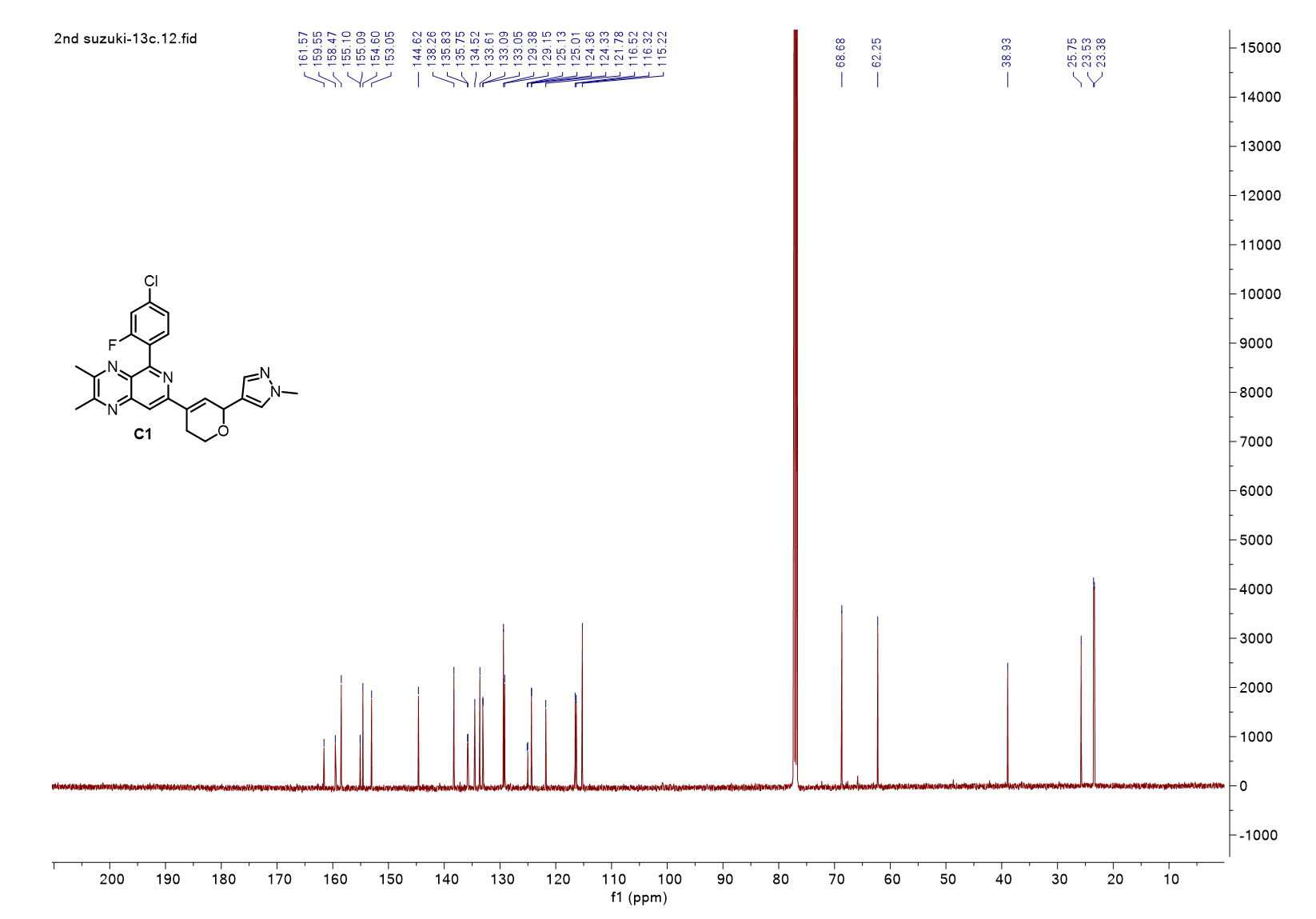

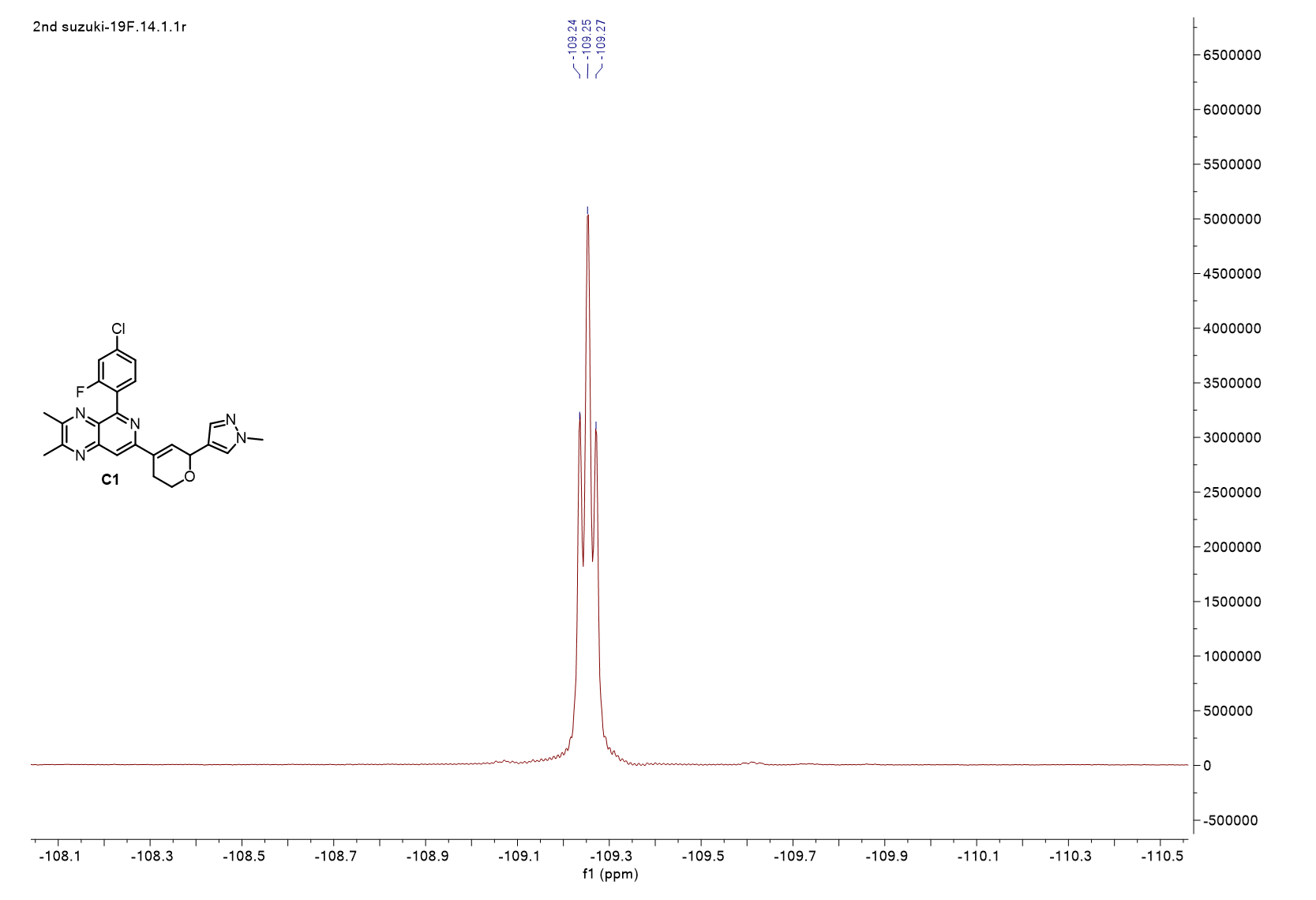

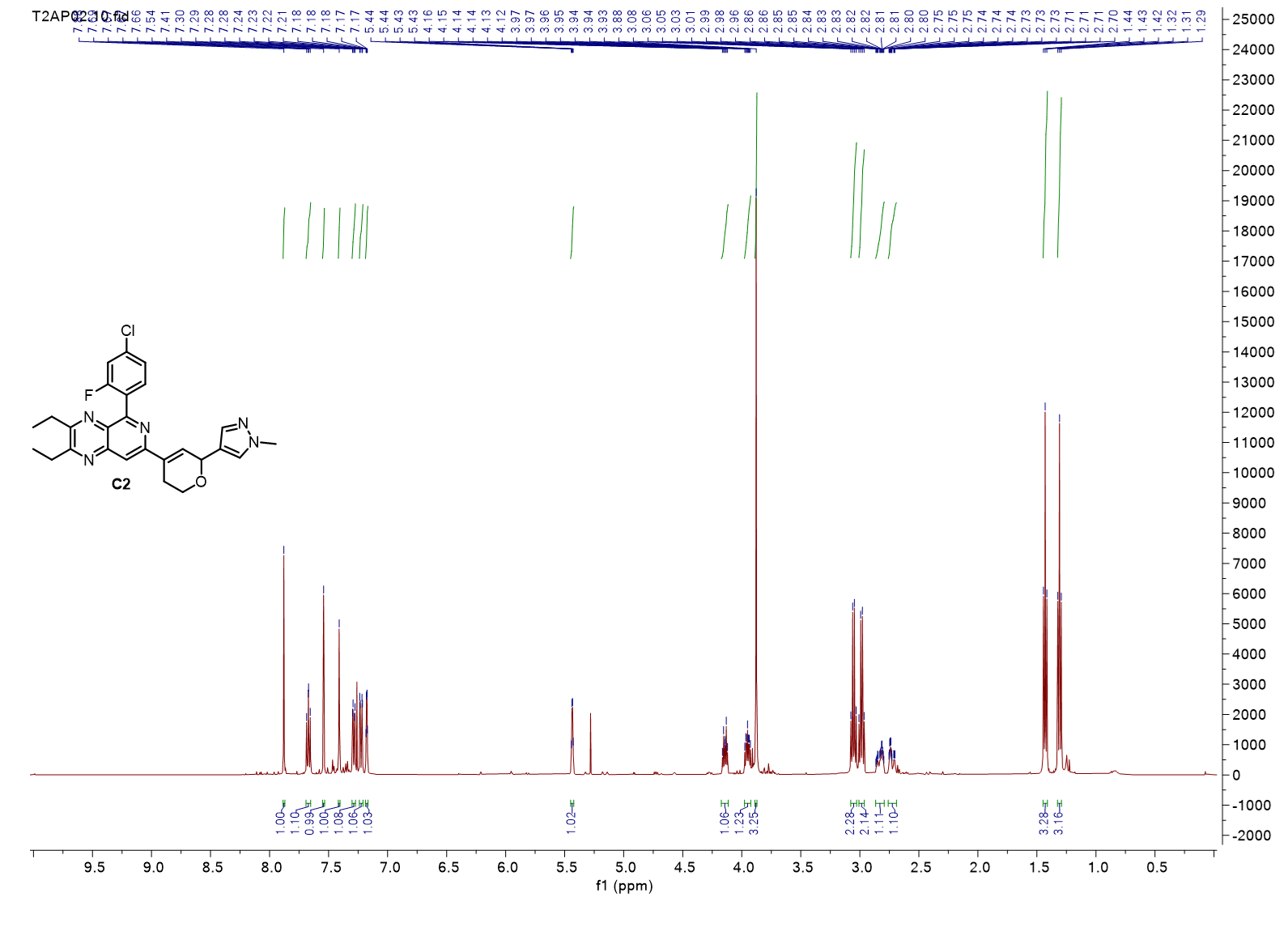

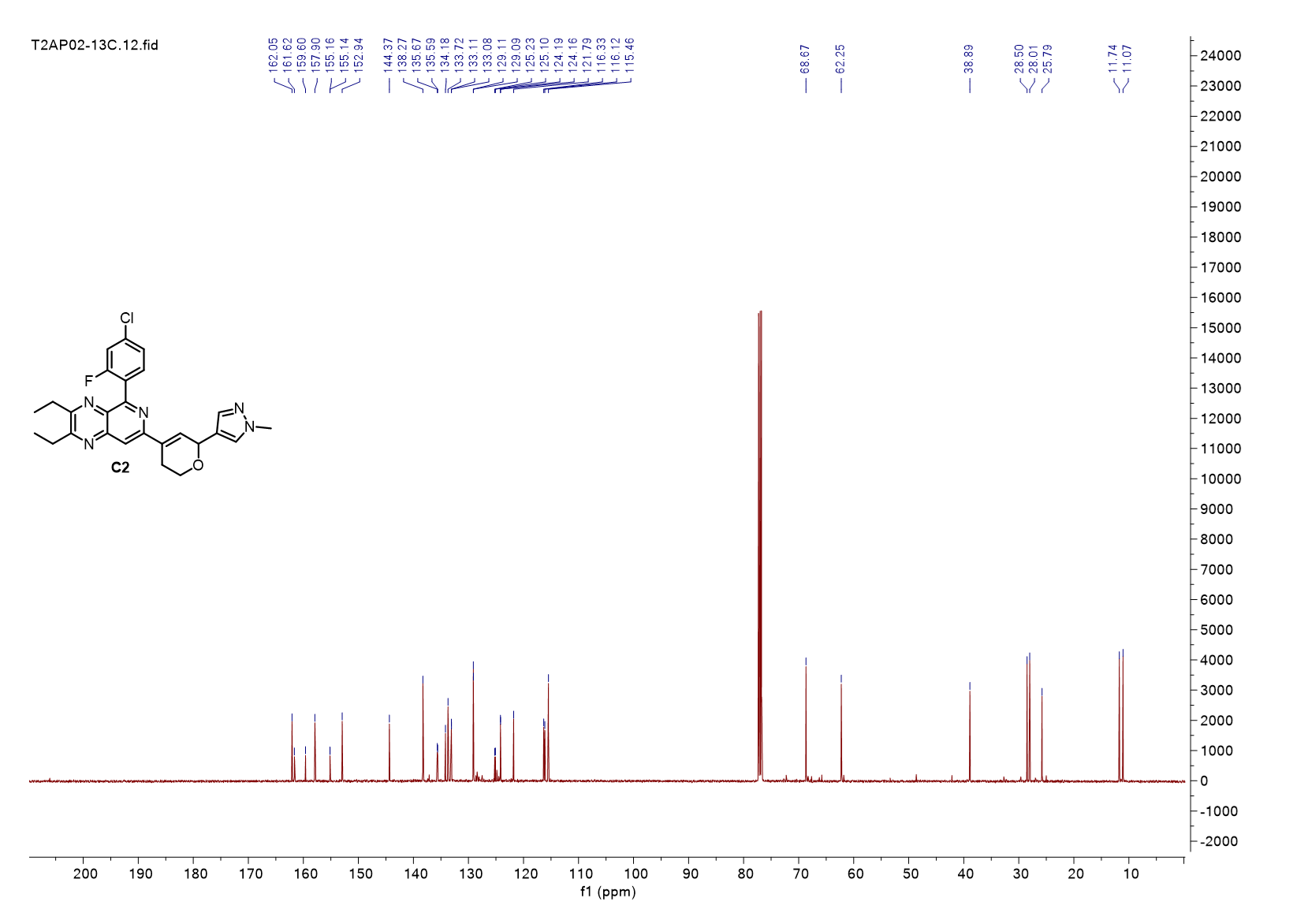

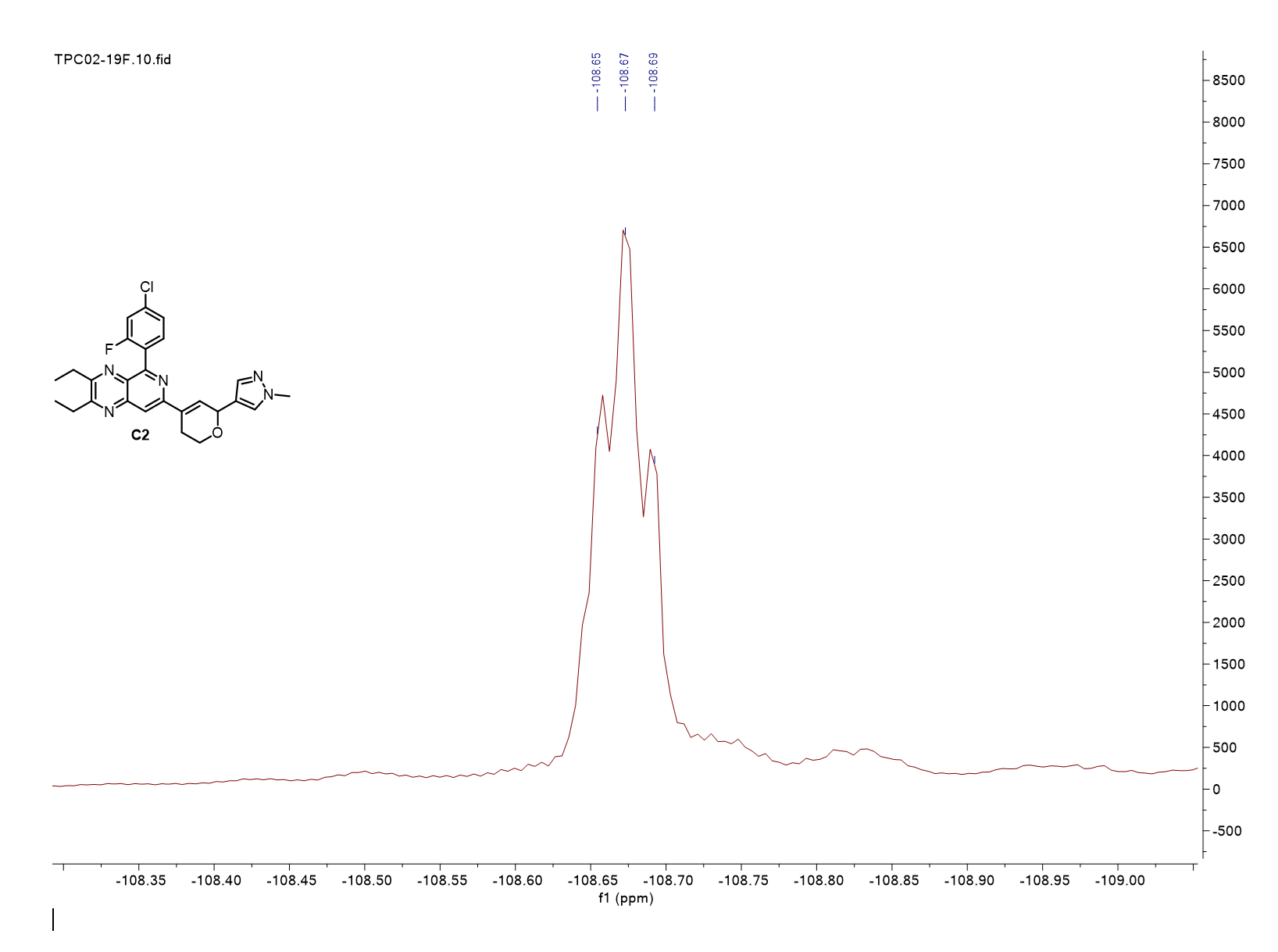

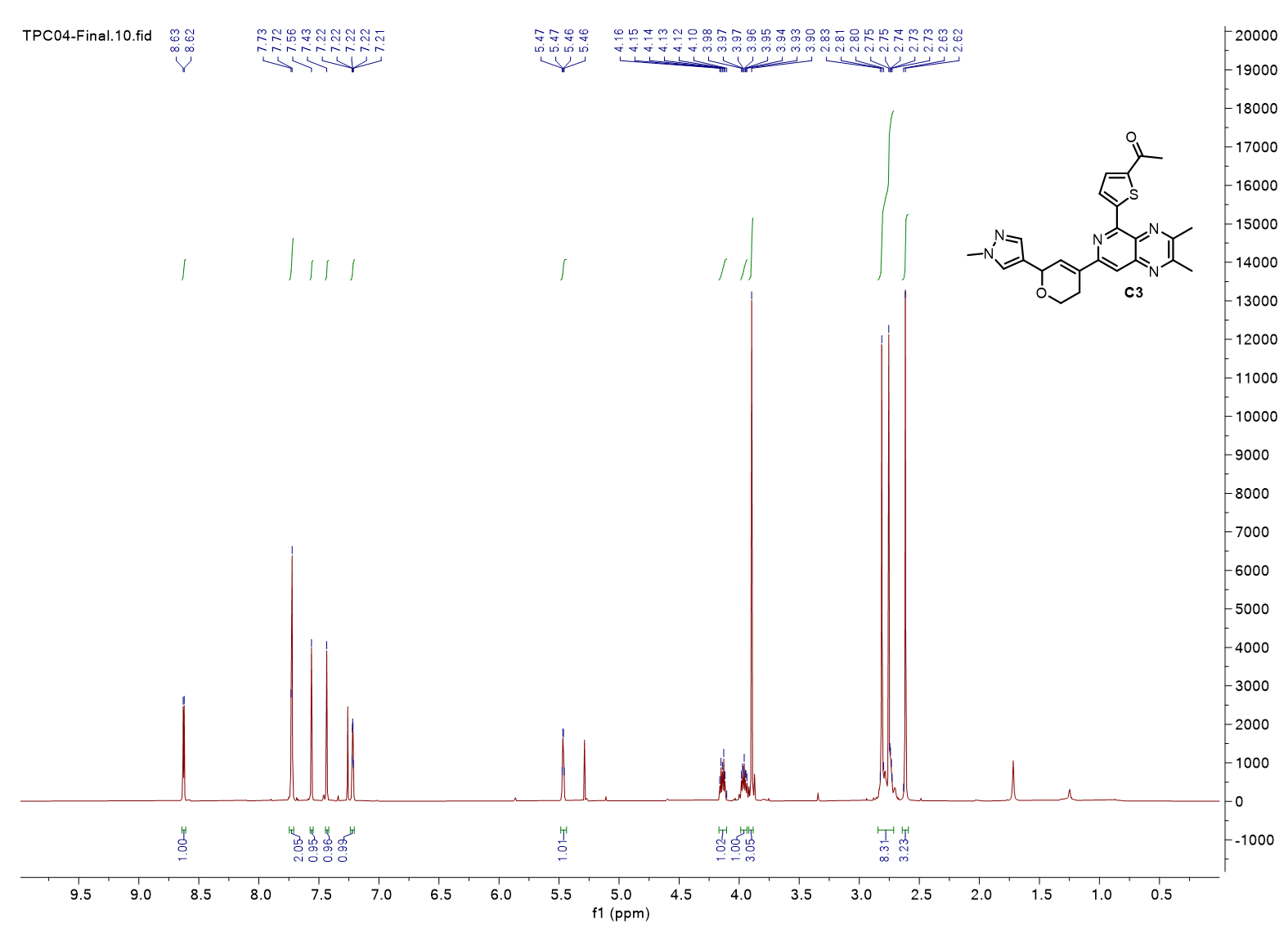

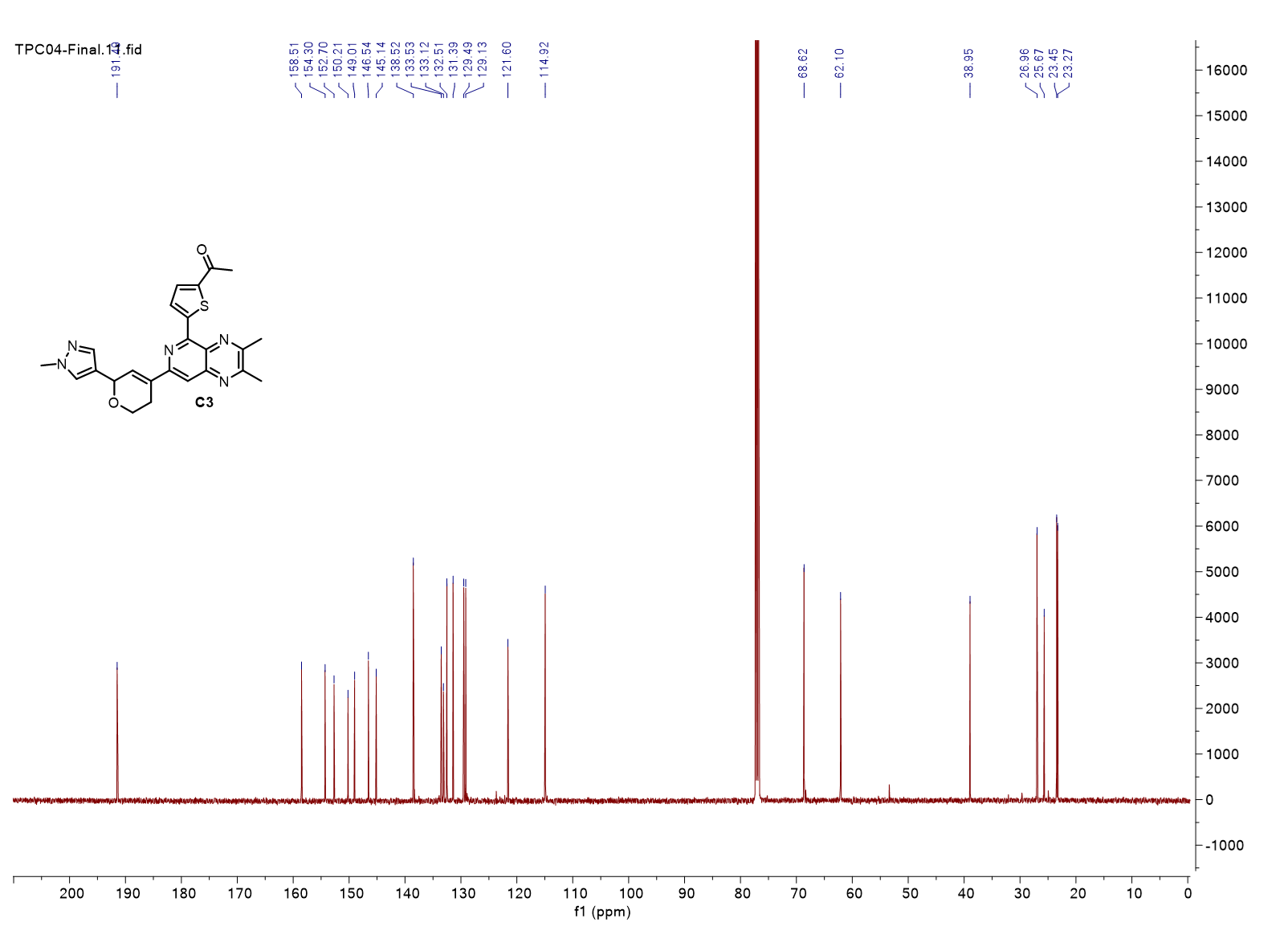

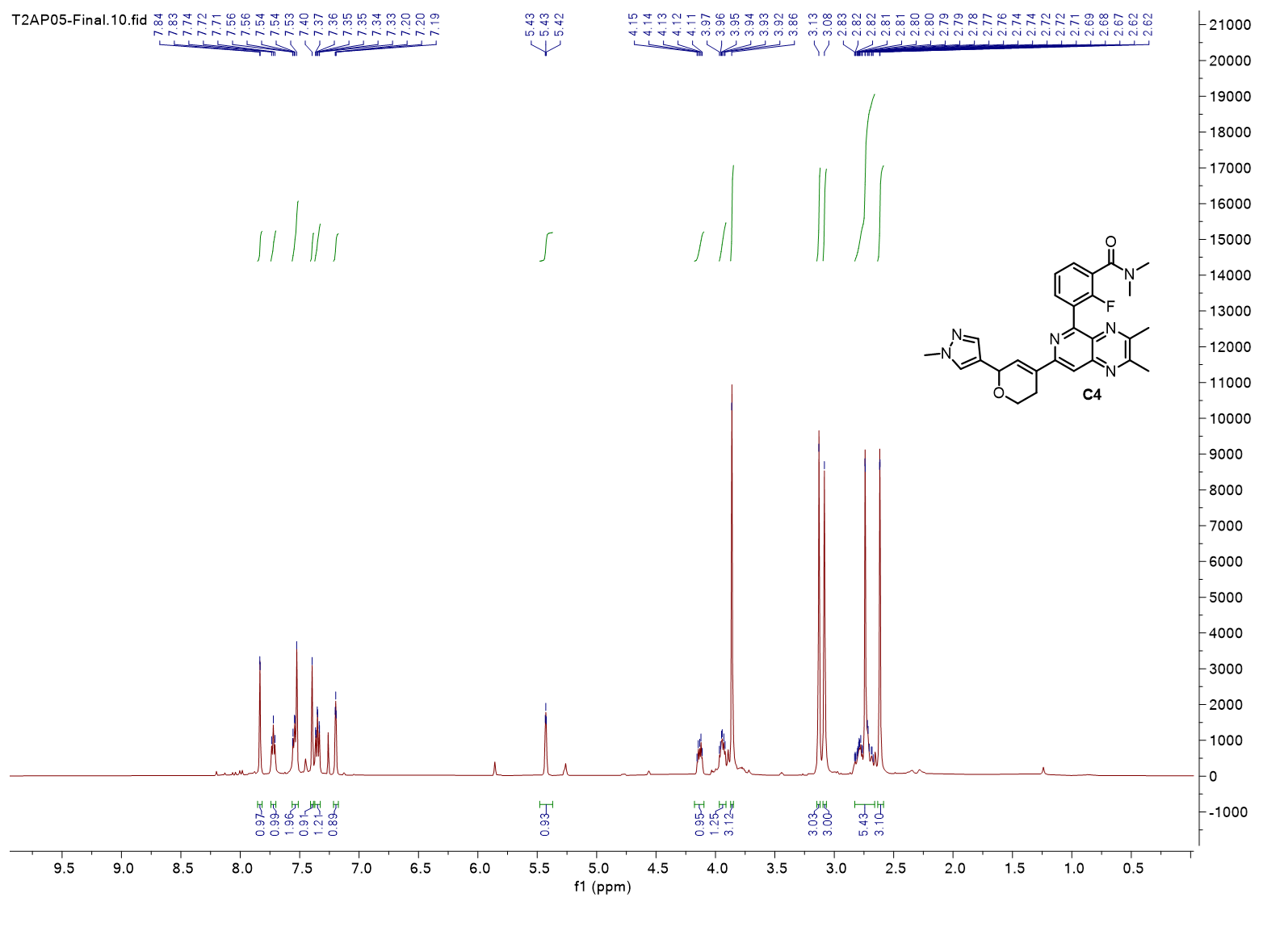

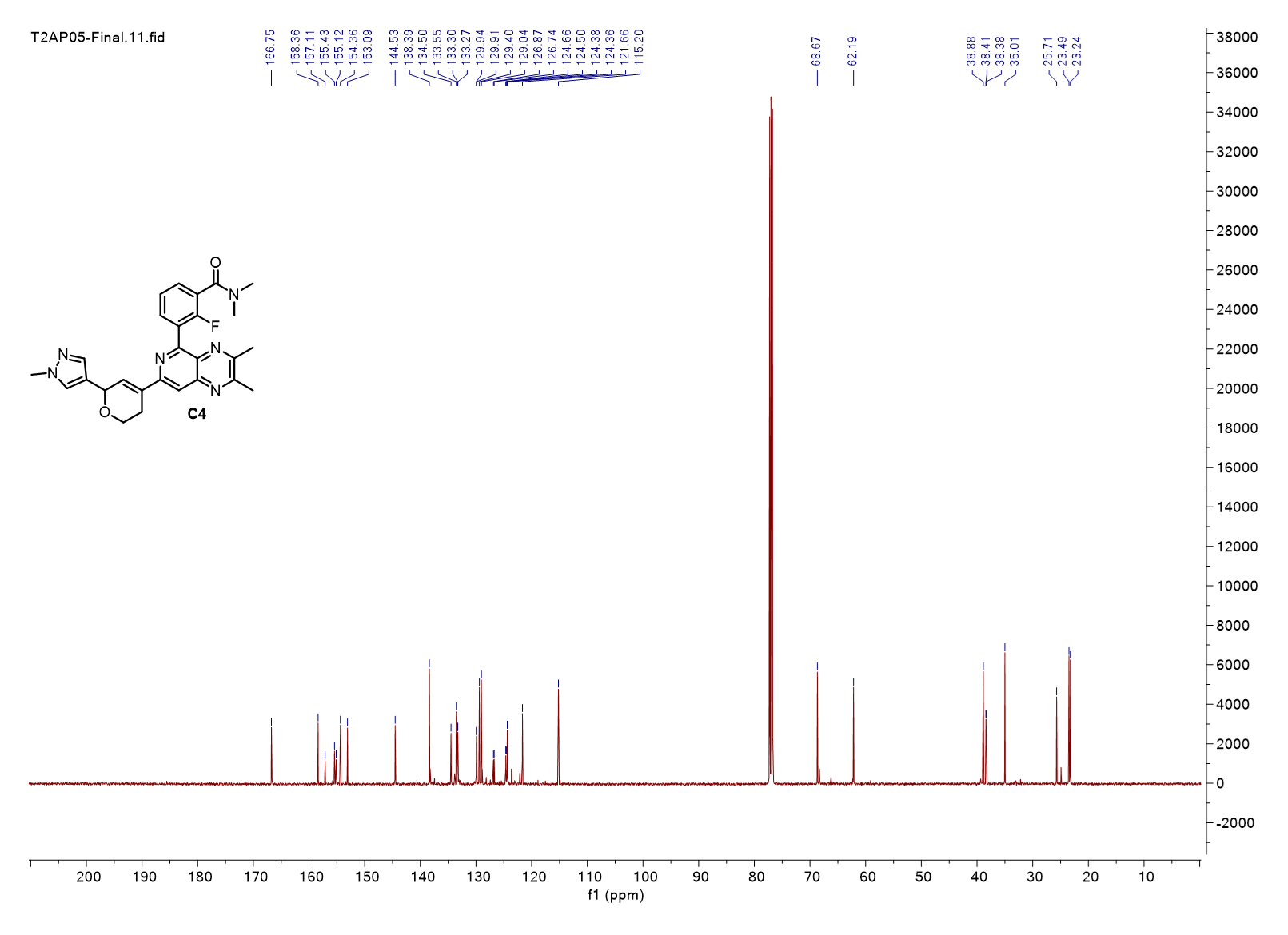

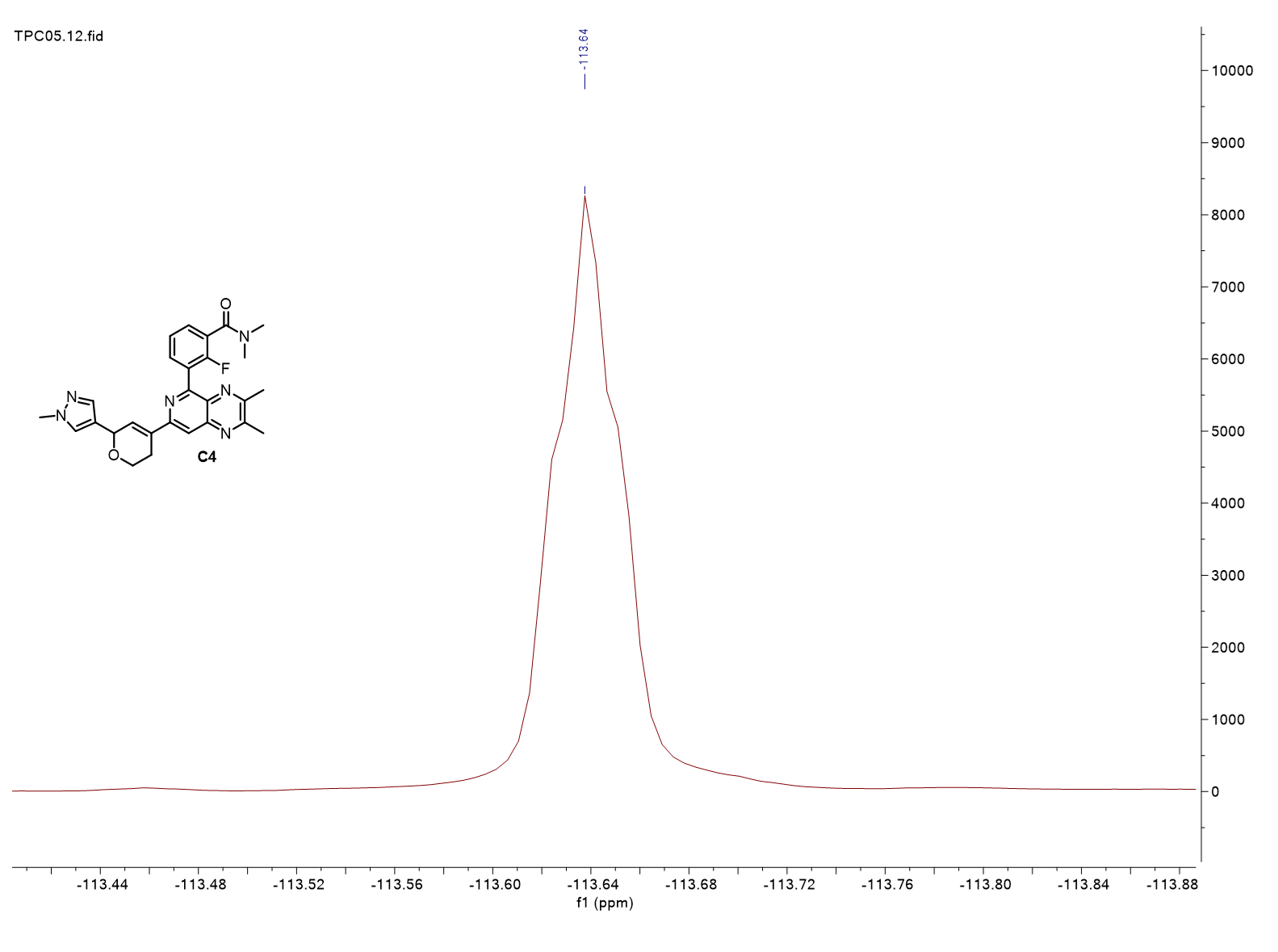

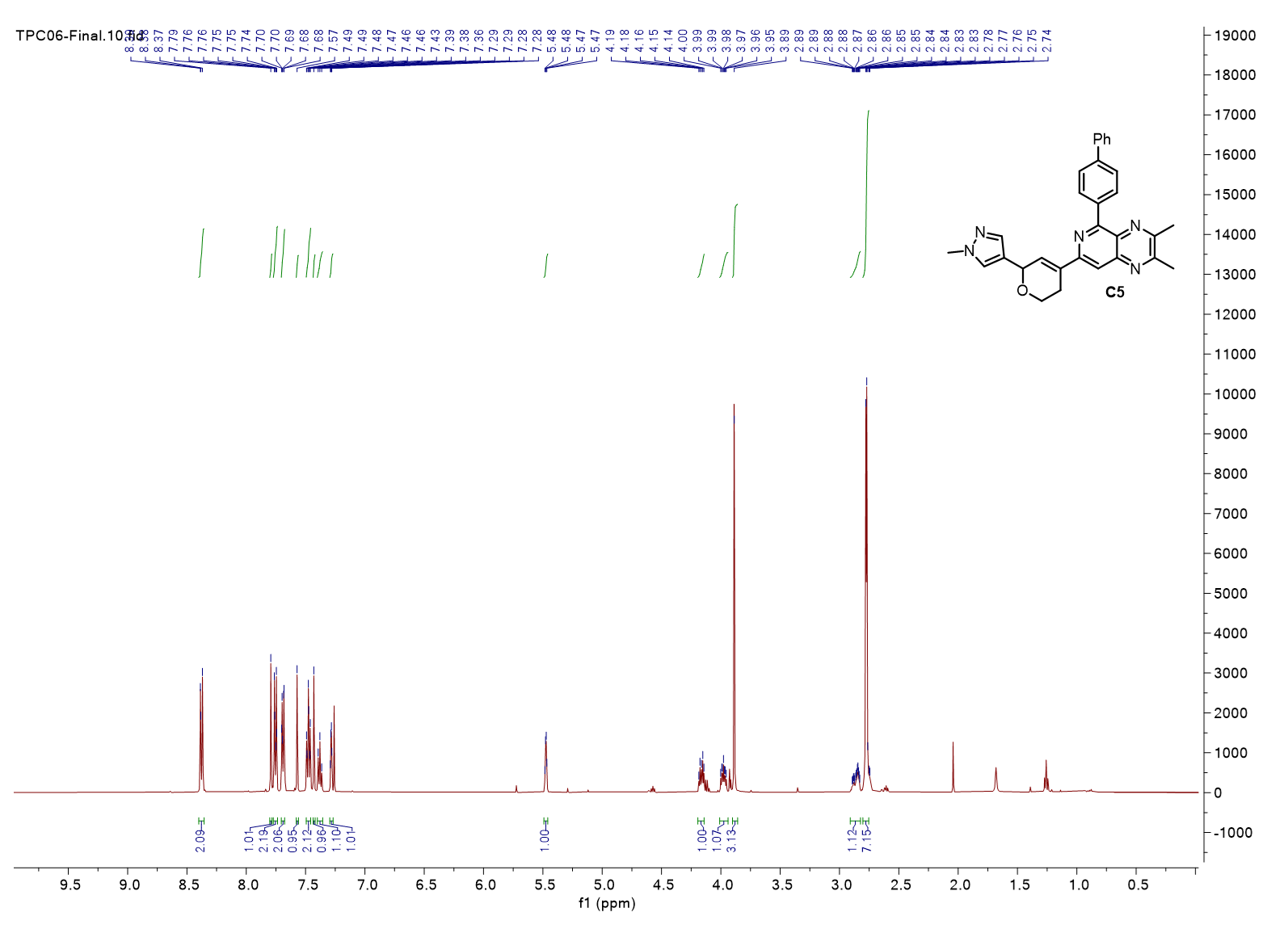

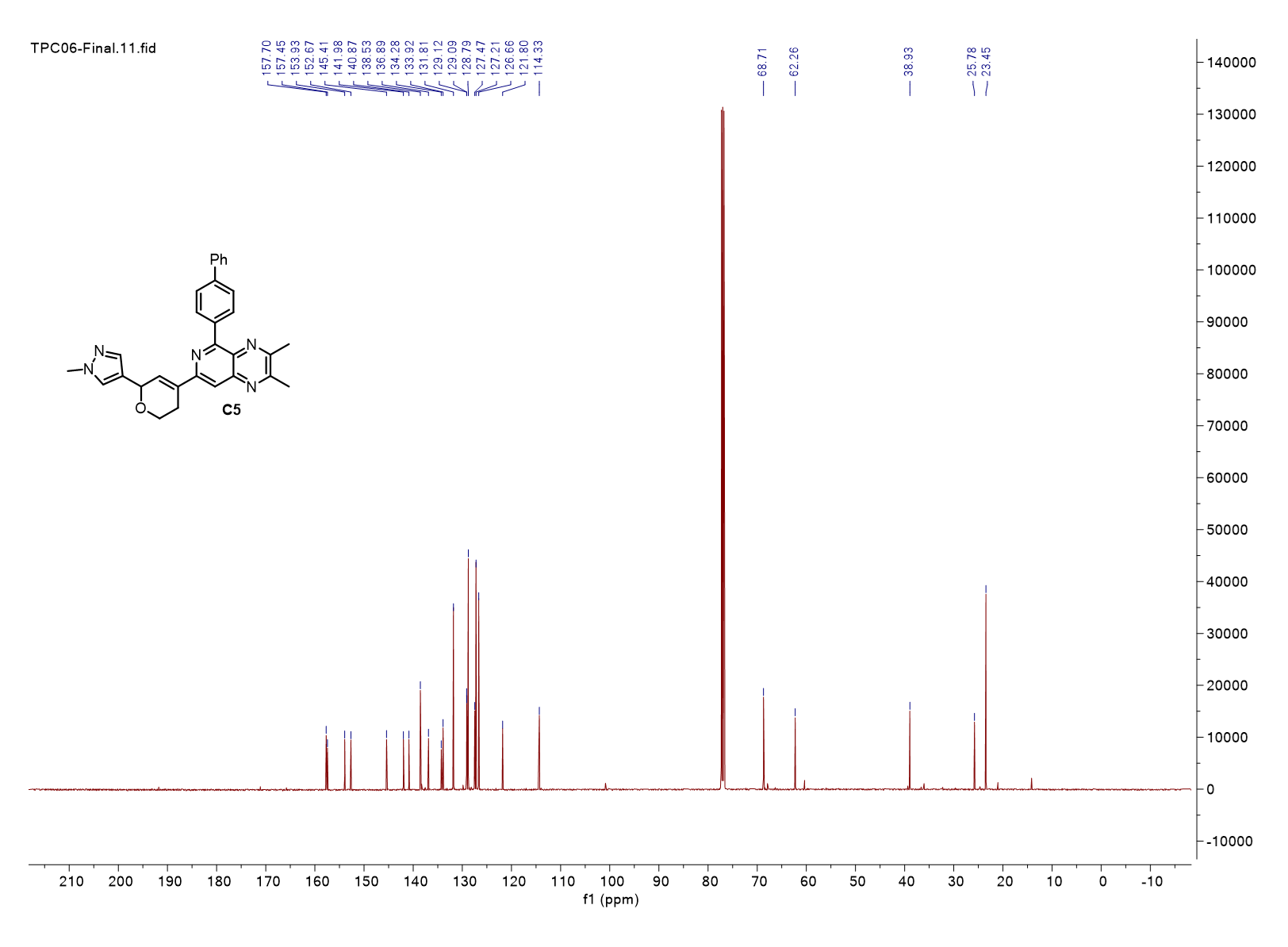

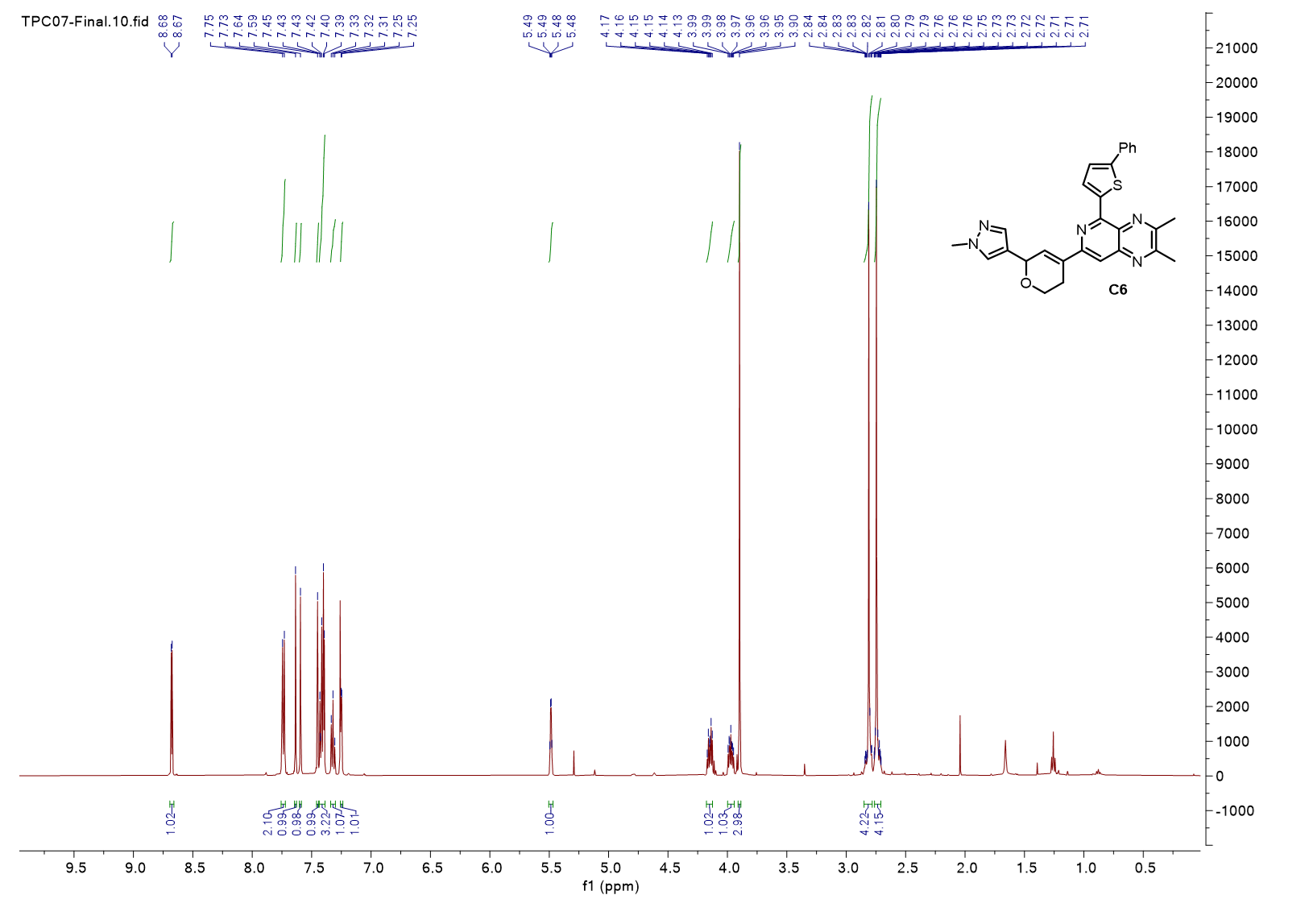

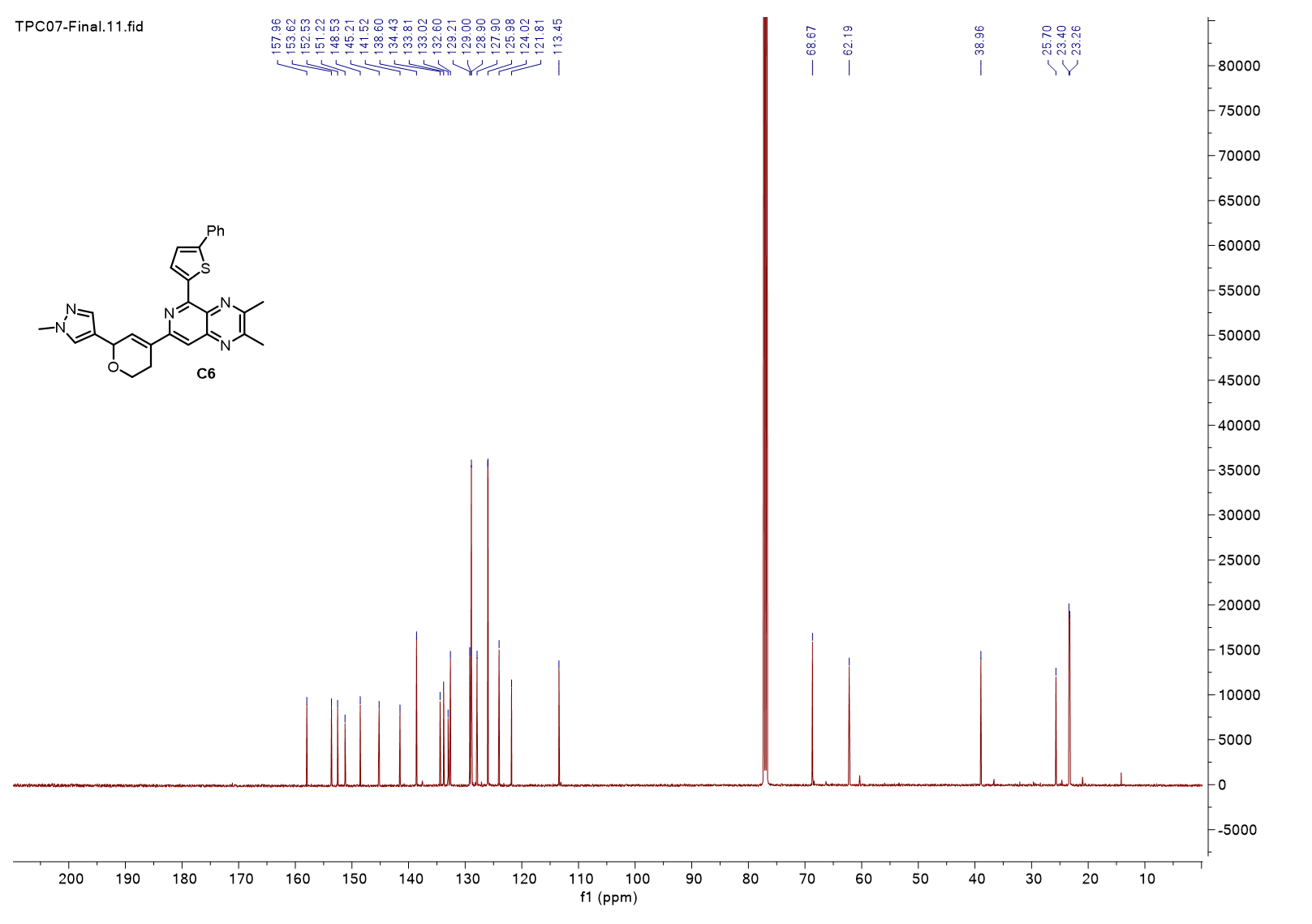

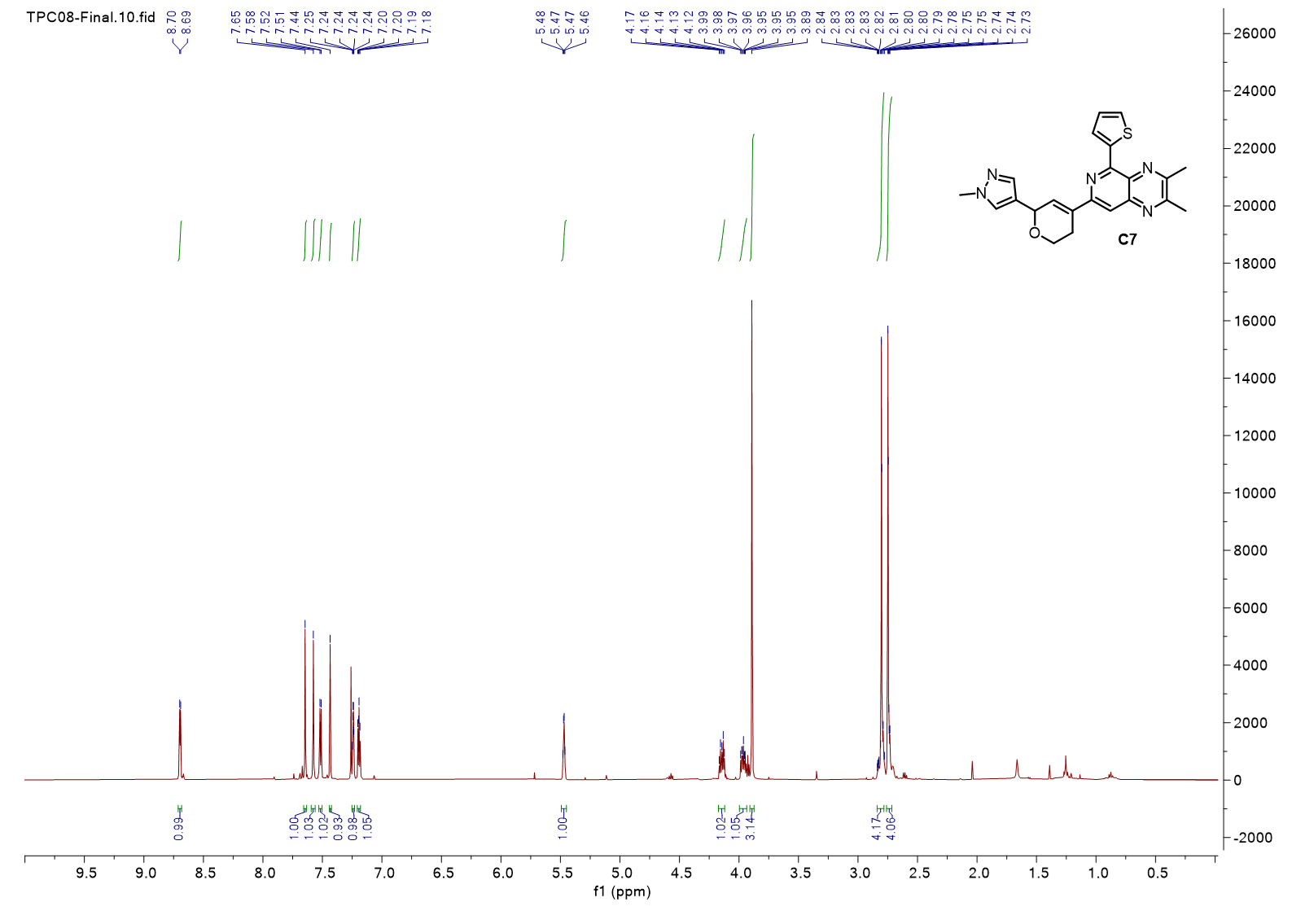

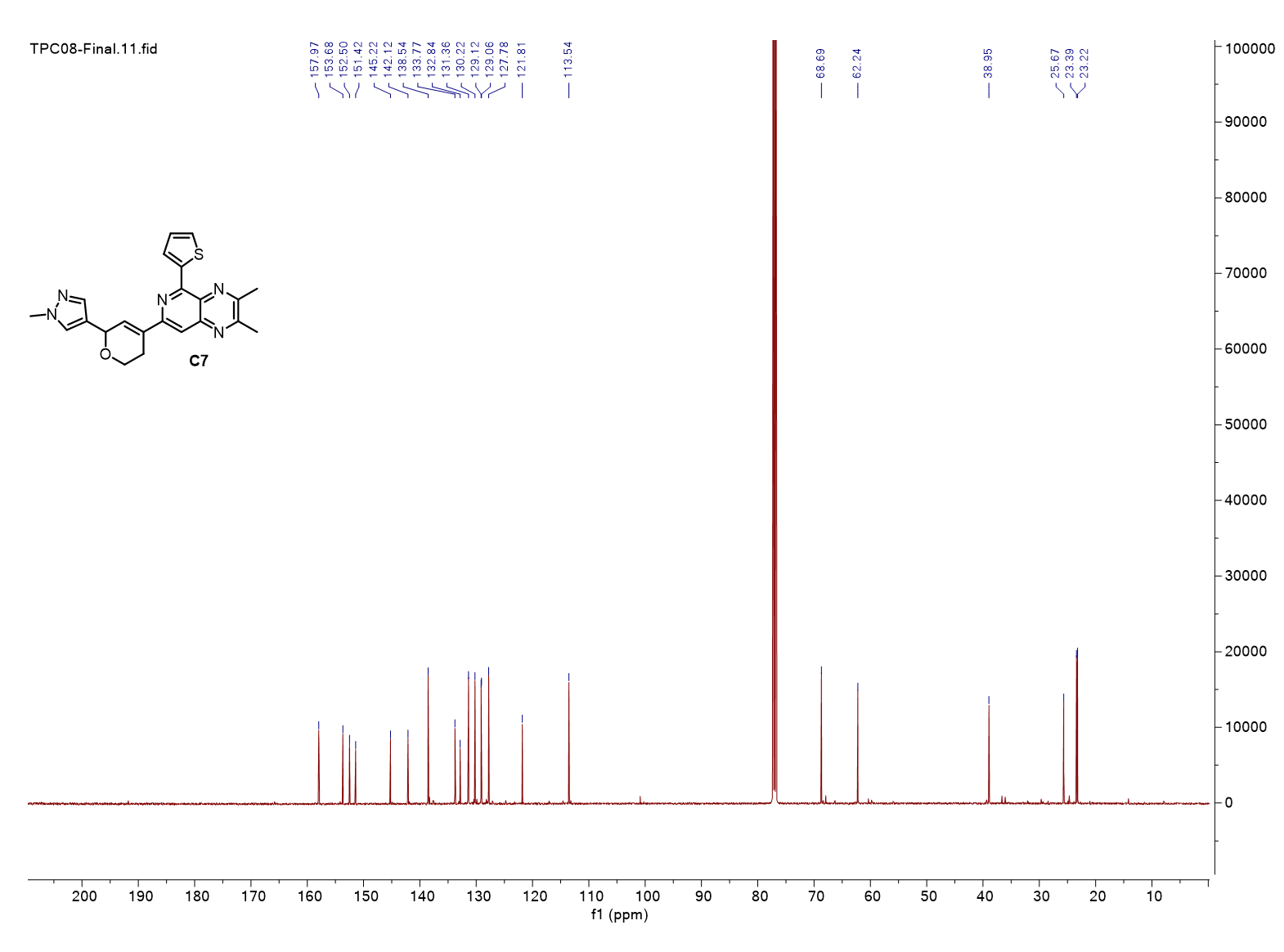

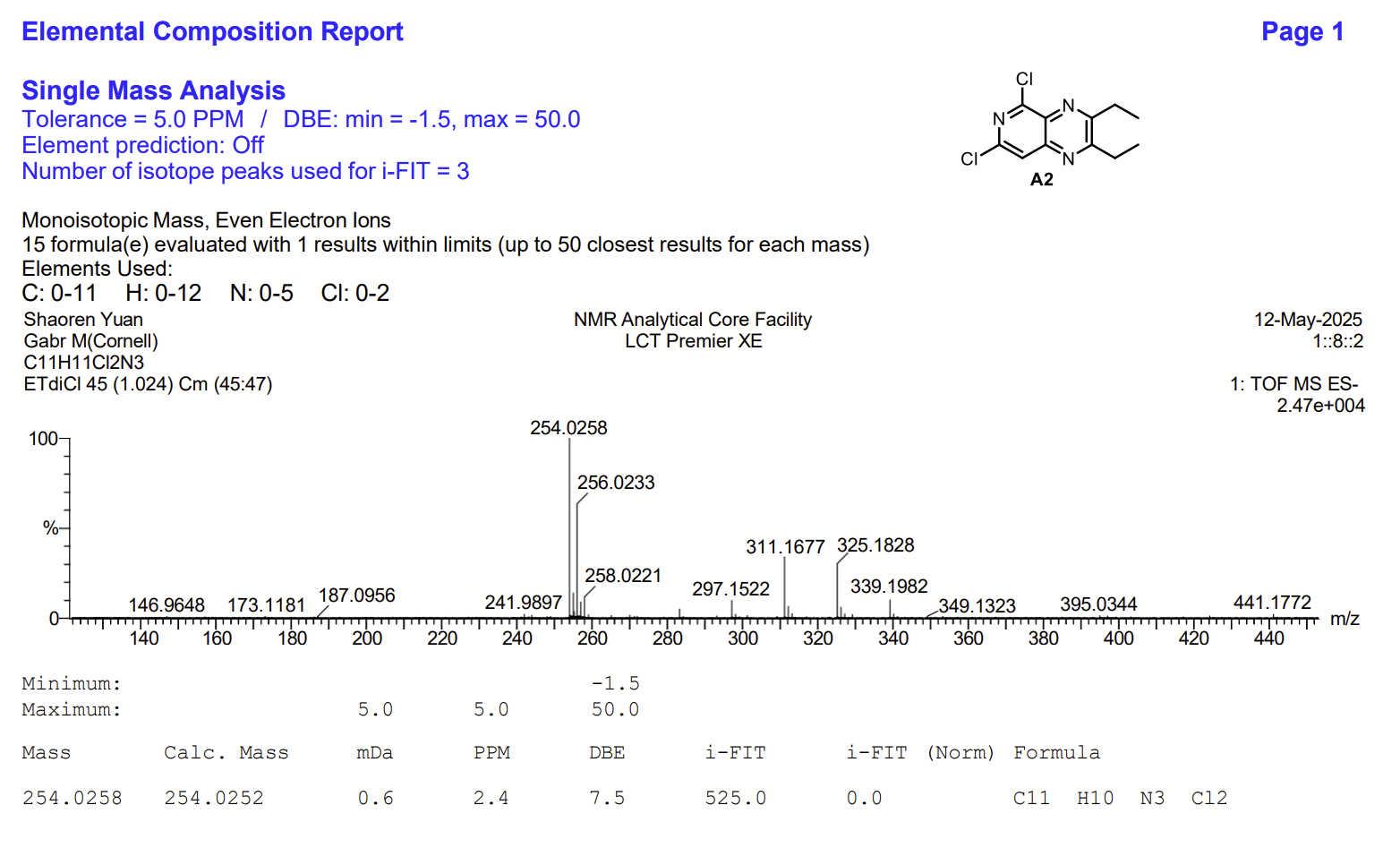

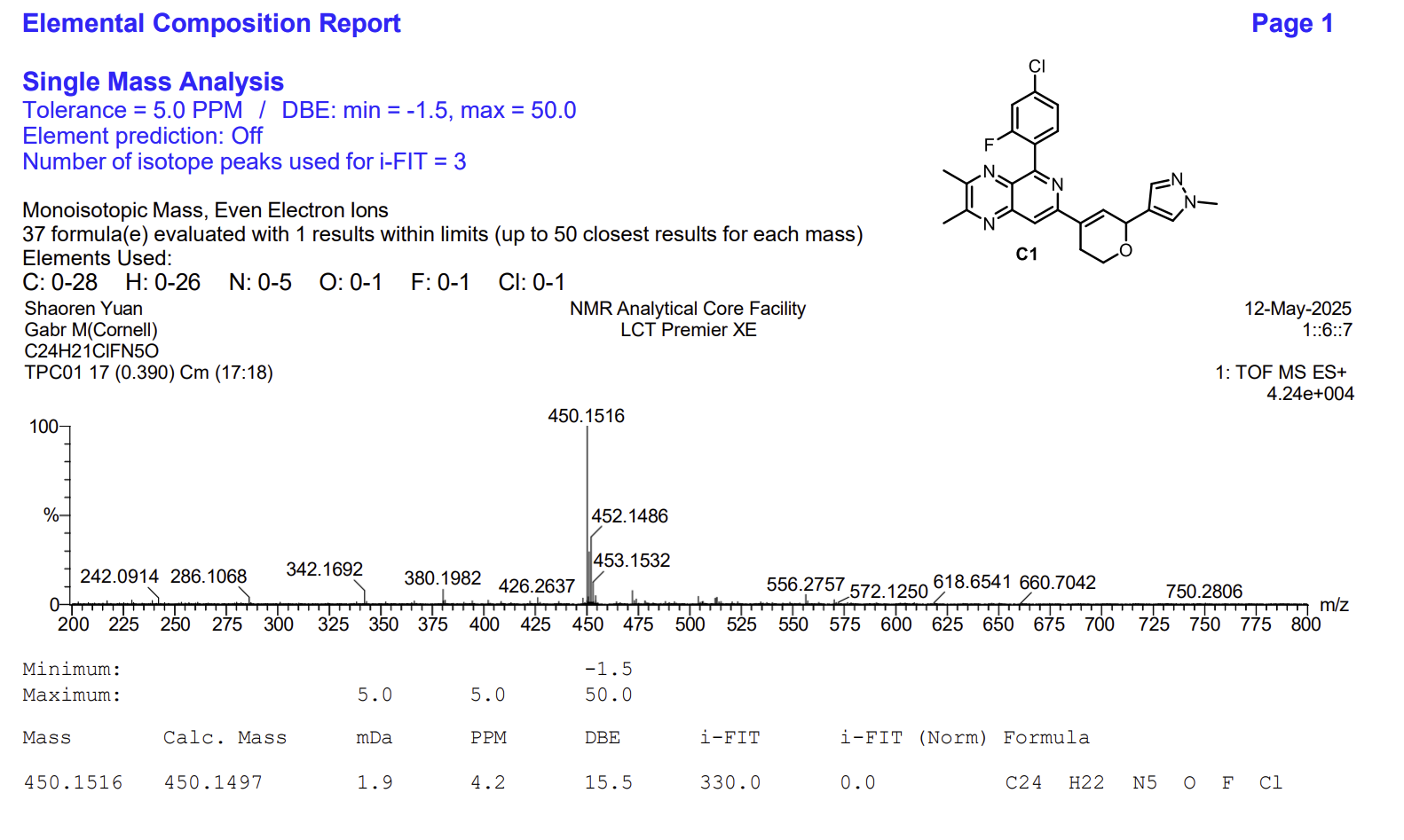

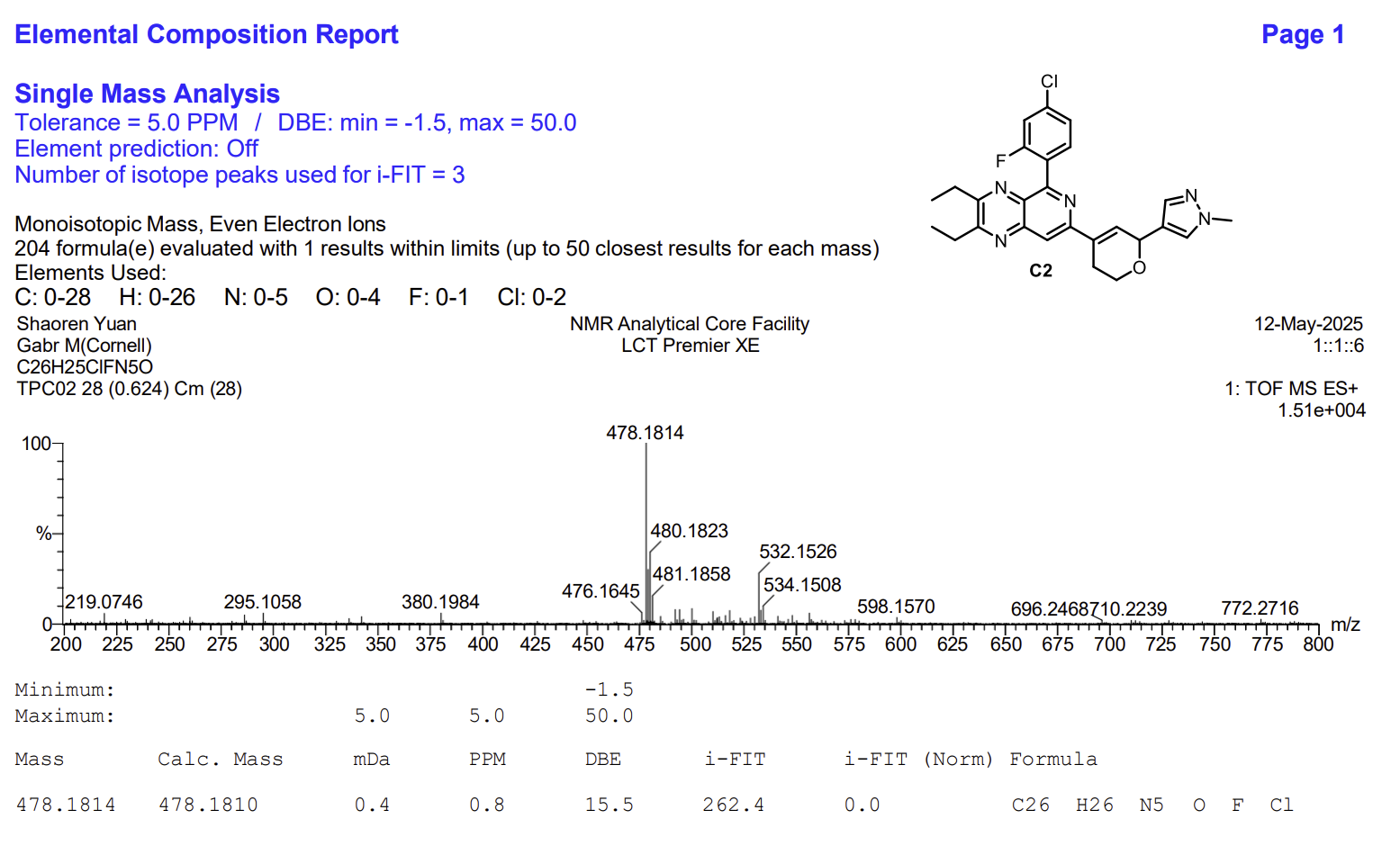

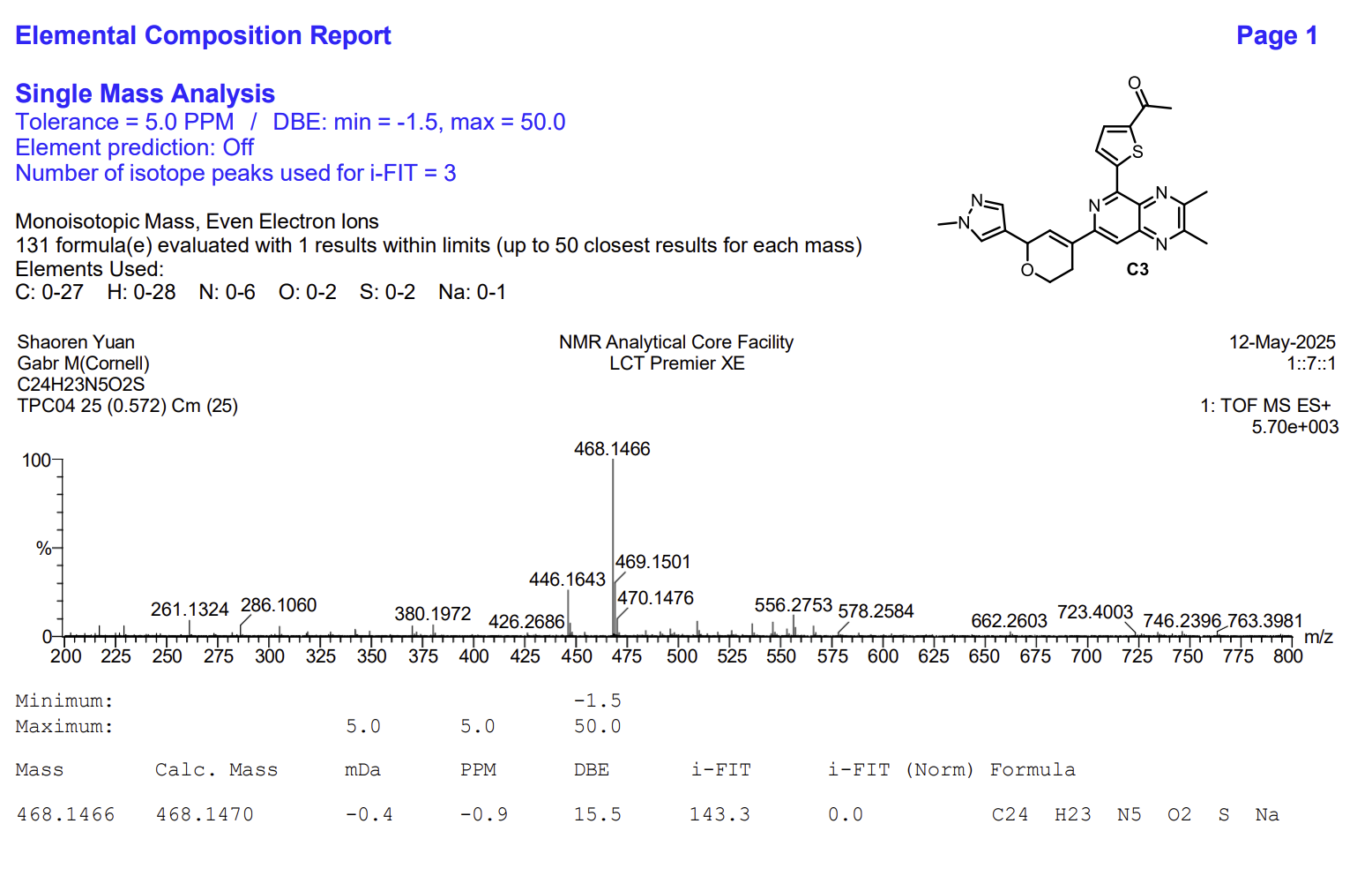

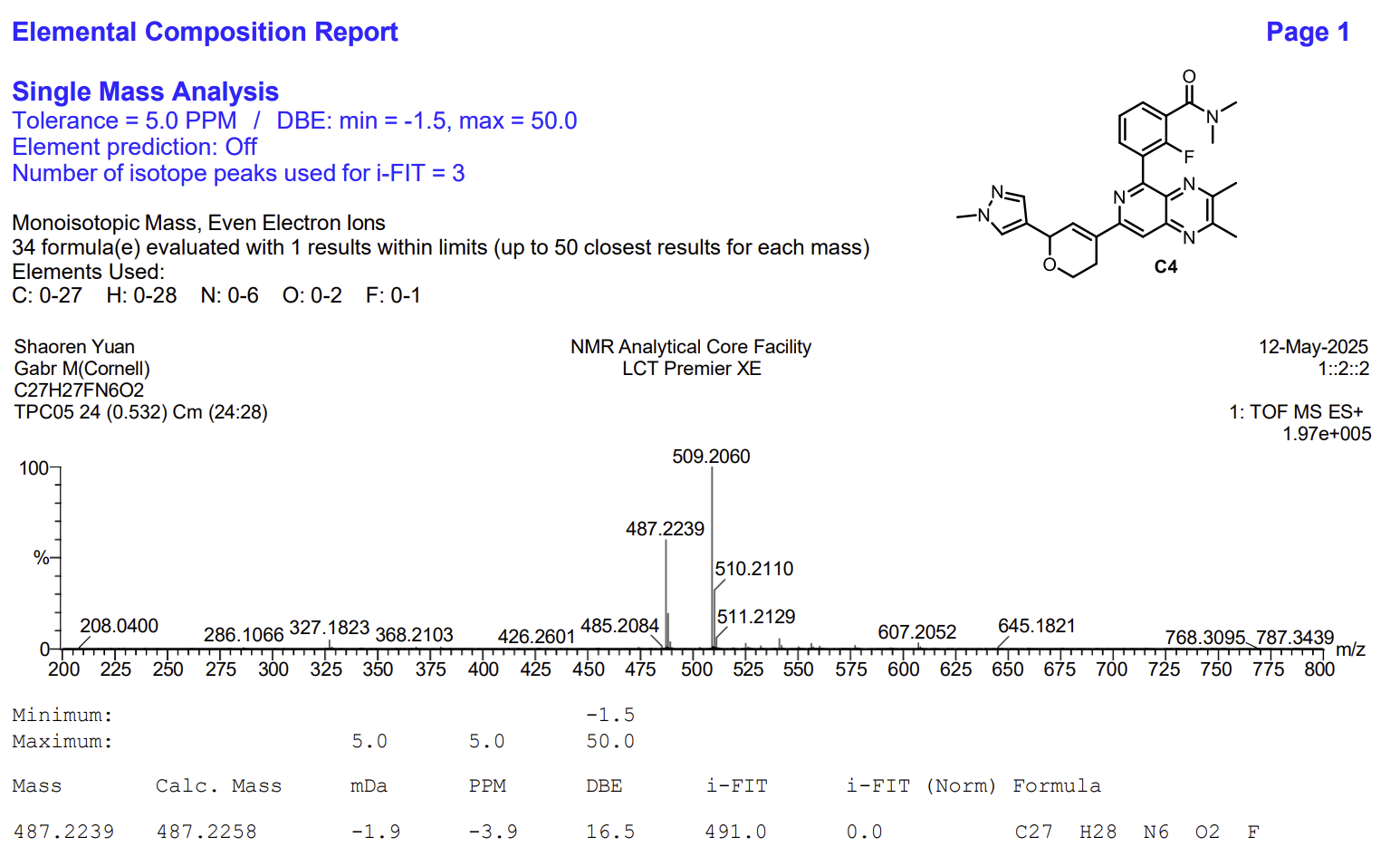

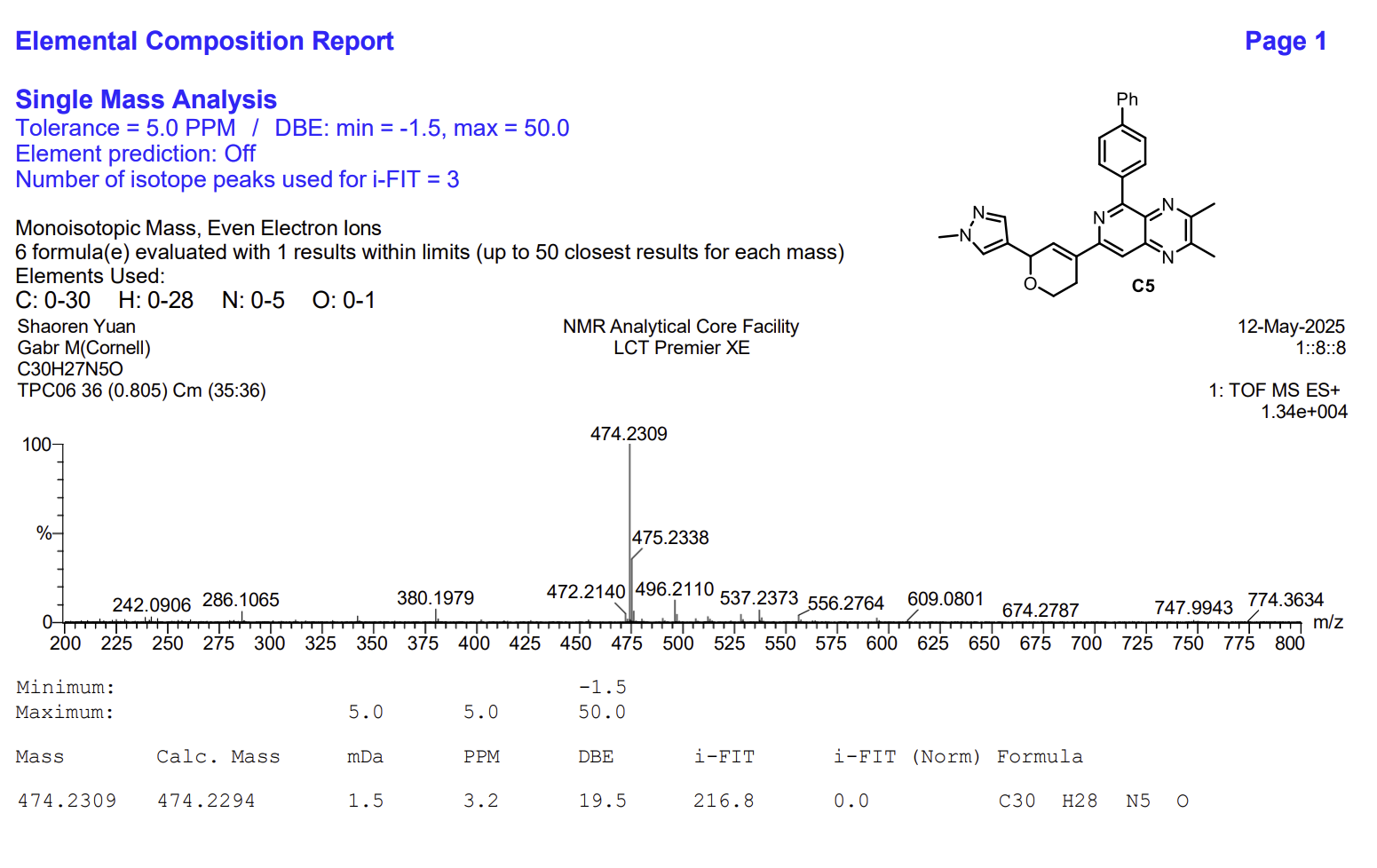

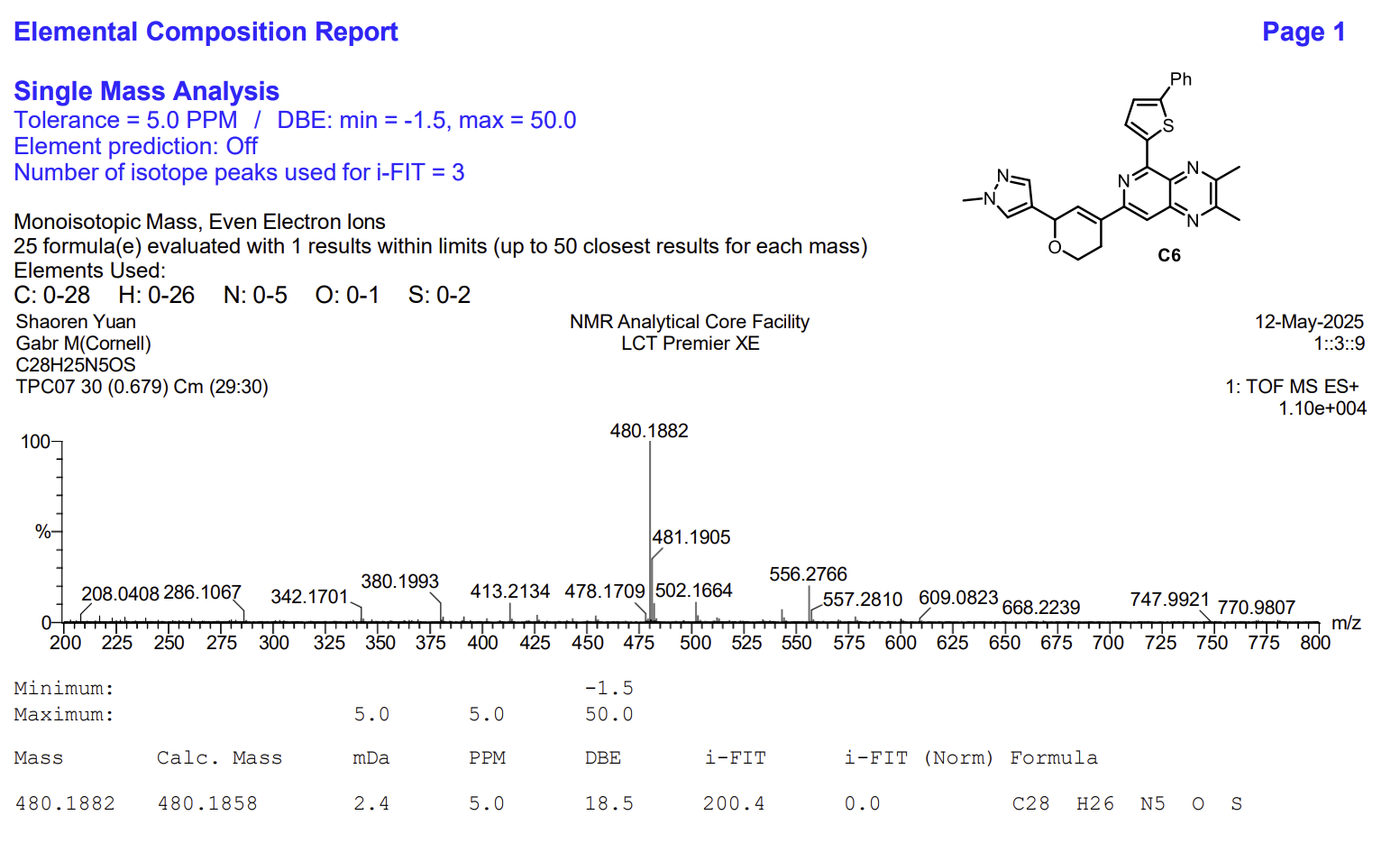

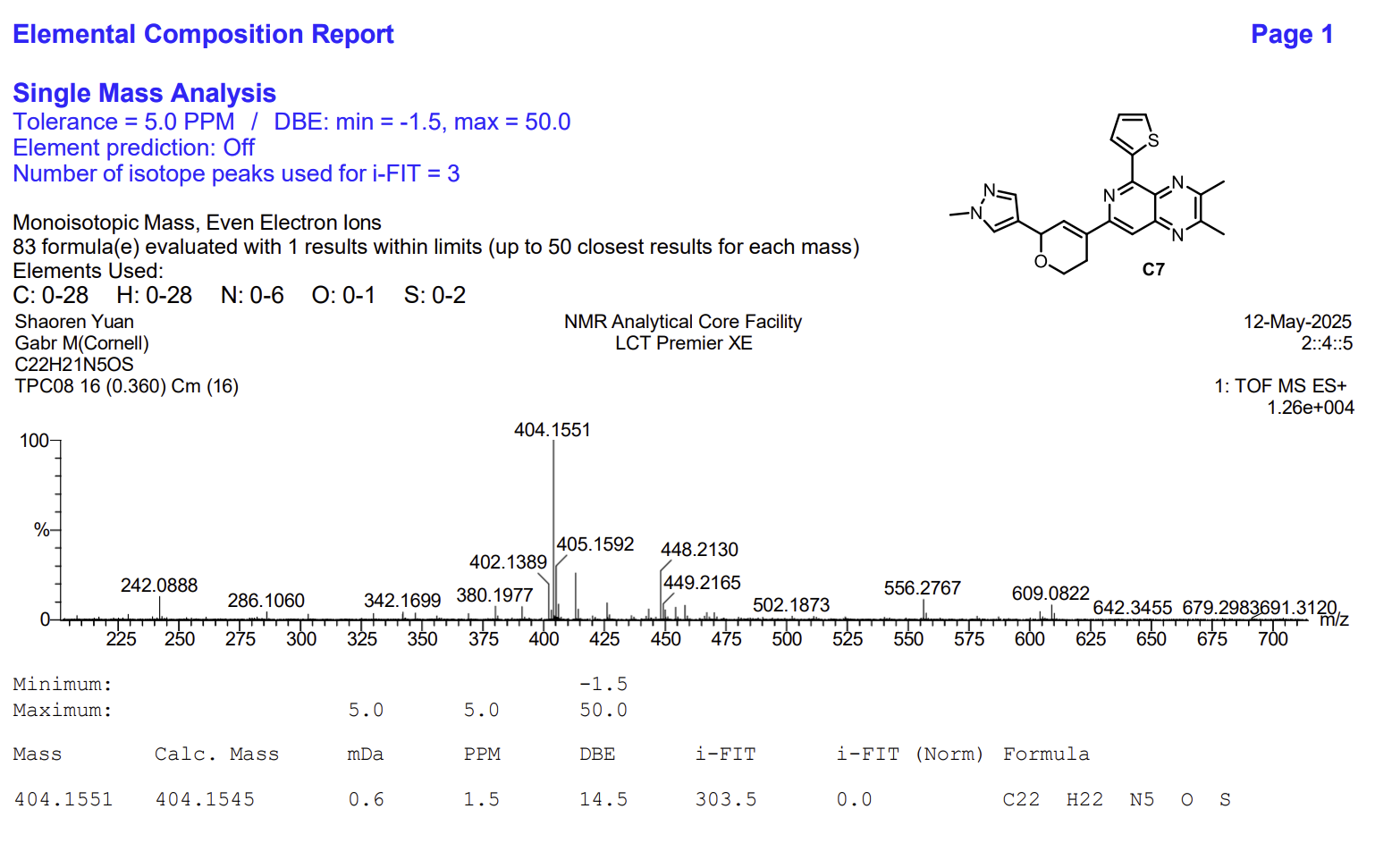
